# Cortical neurons obtained from patient-derived iPSCs with GNAO1 p.G203R variant show altered differentiation and functional properties

**DOI:** 10.1101/2023.06.15.545051

**Authors:** Maria Cristina Benedetti, Tiziano D’Andrea, Alessio Colantoni, Denis Silachev, Valeria de Turris, Zaira Boussadia, Valentina A. Babenko, Egor A. Volovikov, Lilia Belikova, Alexandra N. Bogomazova, Rita Pepponi, Dosh Whye, Elizabeth D. Buttermore, Gian Gaetano Tartaglia, Maria A. Lagarkova, Vladimir L. Katanaev, Ilya Musayev, Simone Martinelli, Sergio Fucile, Alessandro Rosa

## Abstract

Pathogenic variants in the *GNAO1* gene, encoding the alpha subunit of an inhibitory heterotrimeric guanine nucleotide-binding protein (Go) highly expressed in the mammalian brain, have been linked to encephalopathy characterized by different combinations of neurological symptoms, including developmental delay, hypotonia, epilepsy and hyperkinetic movement disorder with life-threatening paroxysmal exacerbations. Currently, there are only symptomatic treatments, and little is known about the pathophysiology of *GNAO1*-related disorders. Here, we report the characterization of a new *in vitro* model system based on patient-derived induced pluripotent stem cells (hiPSCs) carrying the recurrent p.G203R amino acid substitution in Gαo, and a CRISPR-Cas9-genetically corrected isogenic control line. RNA-Seq analysis highlighted aberrant cell fate commitment in neuronal progenitor cells carrying the p.G203R pathogenic variant. Upon differentiation into cortical neurons, patients’ cells showed reduced expression of early neural genes and increased expression of astrocyte markers, as well as premature and defective differentiation processes leading to aberrant formation of neuronal rosettes. Of note, comparable defects in gene expression and in the morphology of neural rosettes were observed in hiPSCs from an unrelated individual harboring the same *GNAO1* variant. Functional characterization showed lower basal intracellular free calcium concentration ([Ca^2+^]^i^), reduced frequency of spontaneous activity, and a smaller response to several neurotransmitters in 40- and 50-days differentiated p.G203R neurons compared to control cells. These findings suggest that the *GNAO1* pathogenic variant causes a neurodevelopmental phenotype characterized by aberrant differentiation of both neuronal and glial populations leading to a significant alteration of neuronal communication and signal transduction.

## INTRODUCTION

The *GNAO1* gene, encoding the alpha-o1 subunit of guanine nucleotide-binding protein (Gαo), has been associated with Developmental and Epileptic Encephalopathies (DEE), a clinically and genetically heterogeneous group of neurological disorders with onset during infancy or childhood (1). Pathogenic variants in the *GNAO1* gene cause encephalopathy affecting psychomotor development with high clinical heterogeneity, including developmental delay, intellectual disability, early-onset hyperkinetic movement disorders with severe exacerbations, epileptic seizures, and prominent hypotonia. Such variants were first described in patients with early infantile epileptic encephalopathy 17 (EIEE17, OMIM #615473) (1). Later, *GNAO1* variants were reported in patients affected by neurodevelopmental disorders with involuntary movements, with or without epilepsy (NEDIM, OMIM #617493) (2). Approximately 50 pathogenic variants have been reported so far in ClinVar (www.ncbi.nlm.nih.gov/clinvar/). Most of them are missense variants in highly conserved residues distributed across the entire length of the protein. Codons 203, 209, and 246 represent pathogenic variant hotspots, occurring in about 50% of cases. A different class of variants leading to *GNAO1* haploinsufficiency has recently been associated with a relatively mild and delayed-onset dystonia phenotype (3;4;5). Recently, the examination of several patients with distinct *GNAO1* variants resulted in a new clinical severity score (6). Interestingly, this study suggested that each variant is associated with a specific severity score and unique disease mechanisms. Despite the increasing number of children diagnosed with *GNAO1*-related disorders, no effective cure or treatments are available.

Gαo is the most abundant membrane protein in the mammalian central nervous system, playing major roles in neurodevelopment (7) and neurotransmission (8). In the brain, Go controls the synthesis of cAMP, modulating inhibitory and stimulatory inputs to the adenylyl cyclase 5 (ADCY5) (9). Go also mediates inhibitory signaling downstream of several neurotransmitters, including dopamine, adenosine, and GABA, through inhibition of calcium channels (10;11) or by preventing neurotransmitter release (12). Go is ubiquitously expressed in the central nervous system, but it is highly enriched in the cerebral cortex, hippocampus, and striatum (13).

Little is known about the molecular mechanism that triggers the neurodevelopmental phenotype in patients carrying pathogenic *GNAO1* variants. Since a link between *GNAO1* variants and rare DEEs was reported for the first time a decade ago (1), several animal models have been raised to understand the pathophysiology of *GNAO1*-related disorders. Knockout models suggest that *GNAO1* plays a pivotal role both in the adult brain and during early neurogenesis. Indeed, its expression increases during axonogenesis (14;15), and knockout mice show impaired neurogenesis (16), while *Gnao1* deficient Drosophila models show defects during axon guidance (17). *GNAO1* disease-linked pathogenic variants in mouse and worm models lead to a loss of function in Gα-mediated signaling with or without a dominant negative effect (18;19;20).

Despite currently available animal models, which include mouse, Drosophila and *C. elegans* (18;19;20;21;22;23;24), greatly contributing to a deeper understanding of this pathology, species-specific differences (e.g. neonatal lethality in p.G203R/+ mice; 22) make it hard to translate findings from animal models to humans. Hence the necessity for more reliable models. In this regard, human induced pluripotent stem cells (hiPSCs) represent a valuable tool for disease modeling and drug screening since they can be derived directly from patients and then differentiated ideally into any cell type of interest. Moreover, gene editing allows for the introduction of specific nucleotide substitutions in existing iPSC lines from healthy individuals, generating pairs of mutated/wild-type isogenic lines. To the best of our knowledge, there is only one hiPSC-based study of *GNAO1*-related disorders (25). The authors showed abnormal structure and aberrant neurite outgrowth in *GNAO1*-KO spheroids (25). They assessed a defect in lamination and found few neural rosettes and the loss of *PAX6* expression compared to wild-type (WT) cells. They confirmed some of these findings in patient hiPSCs carrying the p.G203R substitution. However, a major limitation of this study is the lack of a proper isogenic *GNAO1* WT control for the patient-derived line. Comparison of lines carrying a given pathogenic variant with their isogenic counterpart, indeed, is crucial to distinguish between phenotypes due to the presence of that variant from those originating from different genetic backgrounds (26;27;28).

Here we report the characterization of a new hiPSC-based *in vitro* model of *GNAO1*-related disorders. We have generated cortical neurons from hiPSCs derived from a patient carrying the p.G203R amino acid change and performed molecular, cellular and functional studies at different time points during differentiation. Importantly, all experiments have been carried out in parallel with an isogenic *GNAO1* wild-type line generated by reverting the variant by CRISPR-Cas9-mediated gene editing. Compared to an isogenic control, *GNAO1* p.G203R neurons show altered expression of early and late key differentiation genes, altered timing of differentiation and an increased number of astrocytes, as well as functional impairment. In addition, the altered gene expression and differentiation of neuronal precursor cells was confirmed in a second hiPSC line derived from an unrelated individual harboring the *GNAO1* p.G203R variant.

## RESULTS

### RNA-Seq analysis reveals impaired neuronal fate commitment in *GNAO1* p.G203R neural progenitor cells

HiPSCs derived from an American patient and isogenic control hiPSCs generated by gene editing have been provided by Child’s Cure Genetic Research (see Methods section). In this work, such lines are referred to as GNAO1^+/G203R^ and GNAO1^+/G203G^, respectively (Figure 1A).

**Figure 1.**
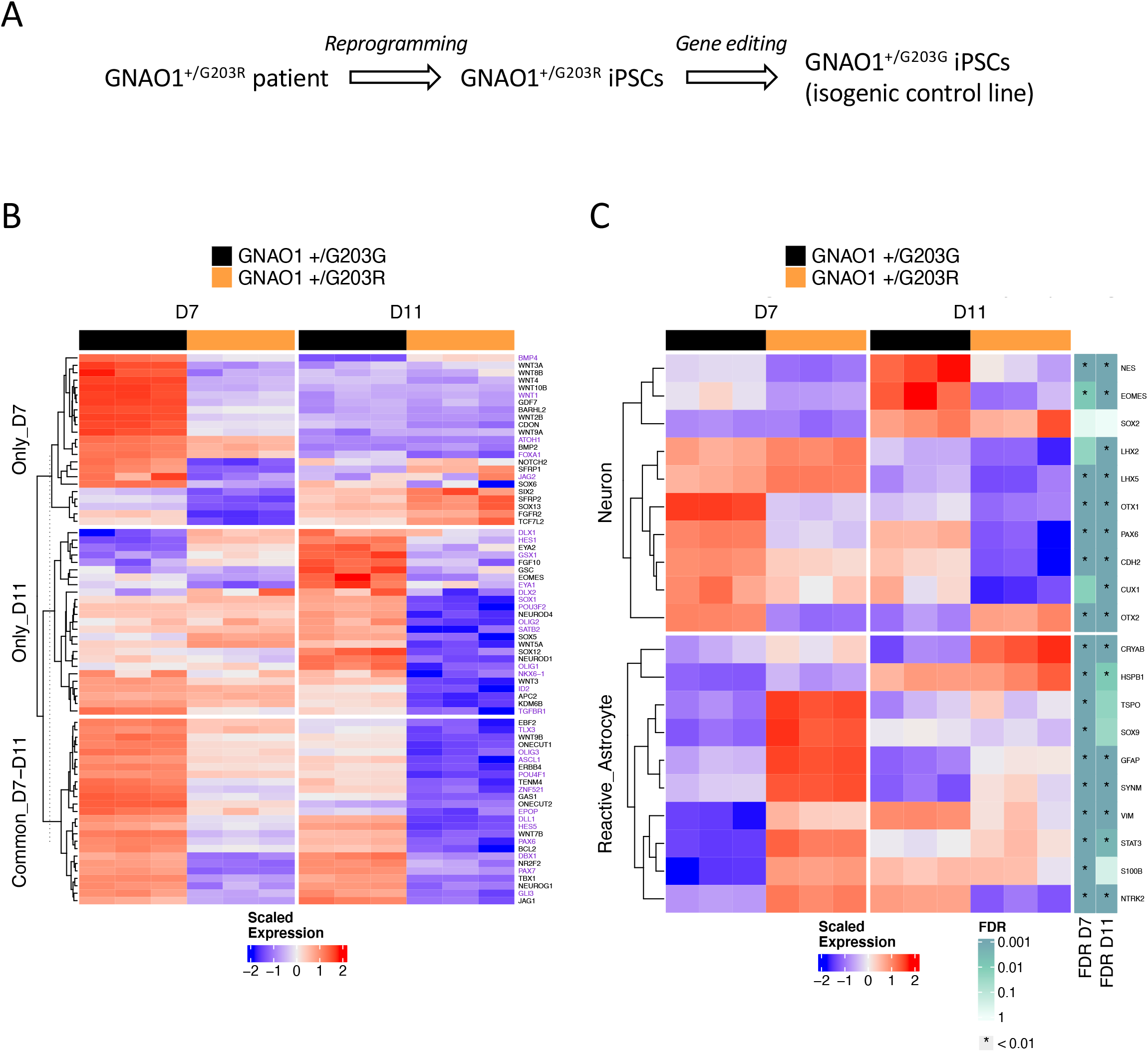
RNA-Seq analysis of neural progenitors. **A.** Outline of the origin of the hiPSC lines used in this study. **B.** Heatmap showing the expression in GNAO1^+/G203R^ and GNAO1^+/G203G^ NPCs at day 7 and 11 of the leading edge genes identified via GSEA for the “cell fate commitment” Gene Ontology Biological Process category. Genes that were identified as leading edge ones only at day 7, only at day 11, or at both time points were split into three groups (Only_D7, Only D11, Common_D7-D11, respectively). Genes belonging to the “neuron fate commitment” or “neuron fate specification” categories were colored in purple. **C.** Heatmap showing the expression in GNAO1^+/G203R^ and GNAO1^+/G203G^ NPCs at day 7 and 11 of selected neuron progenitor- and reactive astrocyte-specific genes. The adjusted p-values displayed on the right pertain to the DGE analyses performed between GNAO1^+/G203R^ and GNAO1^+/G203G^ NPCs at day 7 (FDR D7) and day 11 (FDR D11). The expression values reported in both heatmaps correspond to row-scaled (Z-score), rlog-transformed count data.

We first analyzed RNA-Seq data previously obtained from hiPSC-derived neural progenitor cells (NPCs; day 7 and day 11 of differentiation; n=3 replicates from one differentiation batch). By performing differential gene expression (DGE) analysis, we detected 3,451 downregulated genes and 3,372 upregulated genes at day 7 and 4,174 downregulated genes and 5,081 upregulated genes at day 11 in GNAO1^+/G203R^ NPCs compared to GNAO1^+/G203G^ NPCs (adjusted *p*-value < 0.01) (Supplementary Table S1). These results suggest that at early phases of neural induction the p.G203R variant has an extensive impact on gene expression, as also indicated by principal component analysis (PCA) (Supplementary Figure S1A) showing that the transcriptional profile clearly segregates samples, exhibiting distinct patterns that can be attributed not only to their differentiation stage but also to their genotype. Such effects of the pathogenic variant on gene expression are particularly evident at day 11. We observed concordance between the sets of genes that exhibited deregulation at day 7 and day 11; however, the expression of several genes changed in opposite directions (Supplementary Figure S1B). Gene Set Enrichment Analysis (GSEA) indicated that downregulated genes are enriched in those involved in cell fate commitment, many of which control neuronal differentiation (Figure 1B). Among them, we noticed neural progenitor markers (*PAX6*, *NES/NESTIN*, *EOMES/TBR2*, *CDH2*, *OTX1*) and neuronal markers (*CUX1*), while astrocyte markers were mostly upregulated (Figure 1C). In general, GSEA showed that deregulated genes are involved in many biological processes related to neural differentiation (Supplementary Figure S2A). Specifically, we observed extensive downregulation of genes related to neurodevelopmental processes, such as “forebrain development”, “neuron migration,” “central nervous system”, “neuron differentiation” and “hindbrain development”. We also observed a downregulation of WNT pathway components (29), particularly on day 7, and to a somewhat lesser extent on day 11 (Figure 1B; Supplementary Figure S2B,C). Notably, this pathway regulates neurogenesis and gliogenesis in the cortex with a specific temporal gradient, and downregulation of WNT signaling has been shown to reduce the timing of neurogenesis and trigger precocious astrogenesis (30). Further, Gαo can act as a direct transducer of WNT-Frizzled signaling (31;32), reinforcing the importance of our findings that components of this pathway are downregulated in GNAO1^+/G203R^ NPCs.

Collectively, this RNA-Seq data suggest that during early neurogenesis the *GNAO1* p.G203R pathogenic variant affects cell fate commitment, and in particular neuronal fate, causing the downregulation of genes implicated in neurodevelopment and, on the other hand, the upregulation of astrocyte markers.

### *GNAO1* p.G203R hiPSCs show impaired neuronal differentiation

GNAO1^+/G203R^ hiPSCs, along with the isogenic GNAO1^+/G203G^ control line, were differentiated into cortical neurons using a method previously described (33), based on dual SMAD inhibition followed by the inhibition of SHH signaling with cyclopamine to induce a neural cortical fate (Figure 2A). This protocol recapitulates the steps of neural induction during embryonic development, including the formation of neural rosettes that mimic *in vitro* the forming neural tube. At day 25 of differentiation, when control cells showed the typical radial organization of neural progenitors in rosettes, GNAO1^+/G203R^-derived cells did not acquire such morphology and showed a more advanced phenotype (Figure 2B). Impaired timing of differentiation was confirmed by gene expression analysis performed by qRT-PCR, showing altered levels of neural progenitor markers, such as *NESTIN*, *OTX2* and *PAX6* in patient-derived cells compared to the isogenic control (Figure 2C), in agreement with RNA-Seq analysis at earlier time points (Figure 1B). GNAO1^+/G203R^ cells also expressed lower levels of *TBR2* and *TBR1*, that in the developing neocortex are expressed in intermediate progenitor cells and postmitotic neurons, respectively (34), agreeing with the reduced numbers of TBR1- and TBR2-positive cells in the cortex of mice harboring pathogenic *GNAO1* variants (22). These results were further confirmed by immunostaining analysis at the same time point, showing a reduced number of PAX6-, TBR2- and NESTIN-positive progenitors in patient-derived cells compared to the isogenic control (Figure 3).

**Figure 2.**
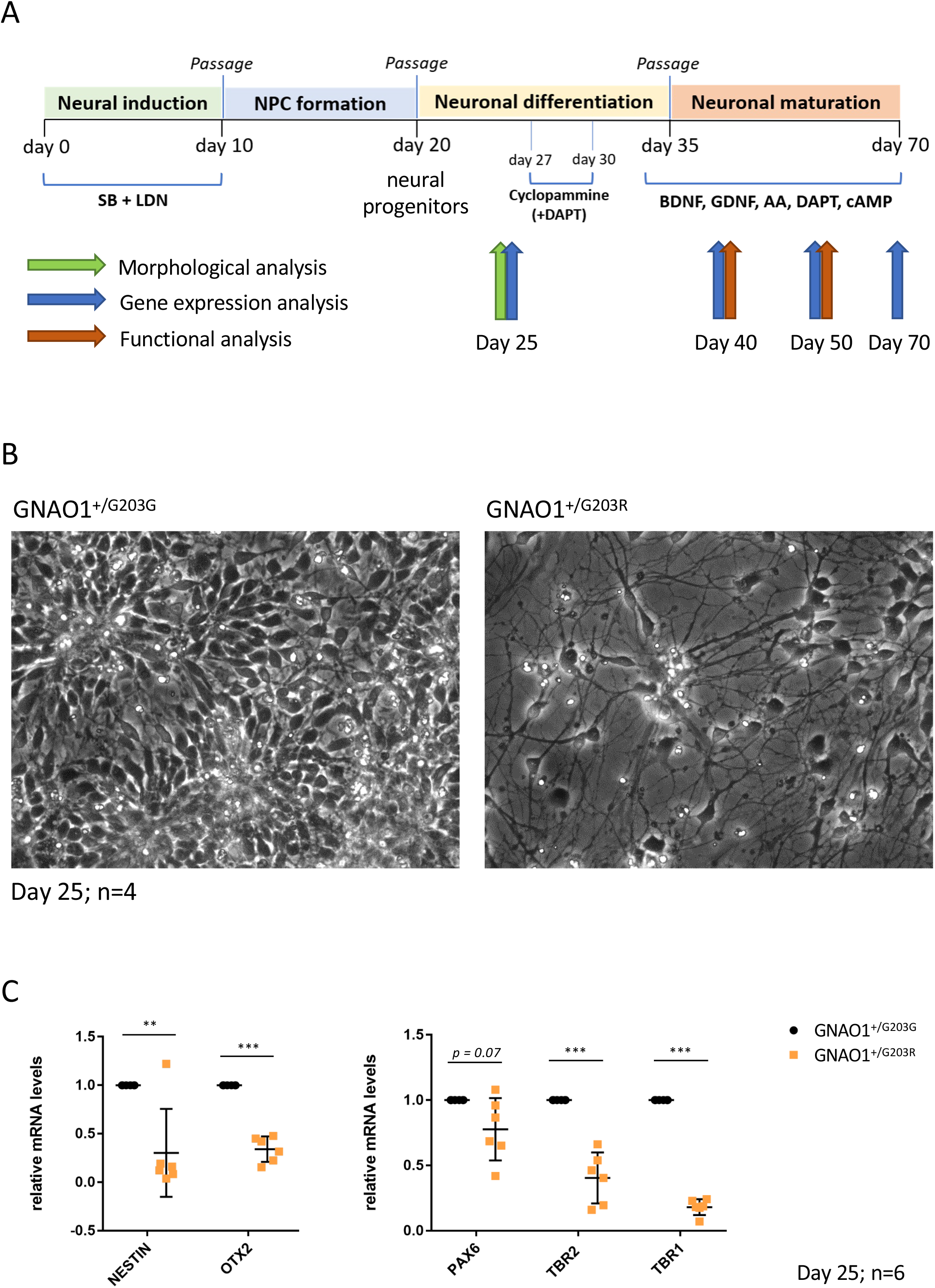
Cortical neuron differentiation and analysis at day 25. **A.** Schematic representation of the protocol used for cortical neuron differentiation of human iPSCs in conventional 2D cultures. NPC: neural progenitor cell; SB: SB431542 (TGFβ type I receptor/ALK5 inhibitor); LDN: LDN193189 (BMP inhibitor); DAPT: Notch inhibitor; BDNF: Brain Derived Neurotrophic Factor; GDNF: Glial Derived Neurotrophic Factor; AA: ascorbic acid; cAMP: cyclic AMP. **B.** Phase contrast images of differentiating cells at day 25. **C.** Real-time qRT-PCR analysis of the expression of the indicated markers in differentiating cells. The graphs show the average and standard deviation (Student’s t test; paired; two tails ; **p<0.01, ***p<0.001).

**Figure 3.**
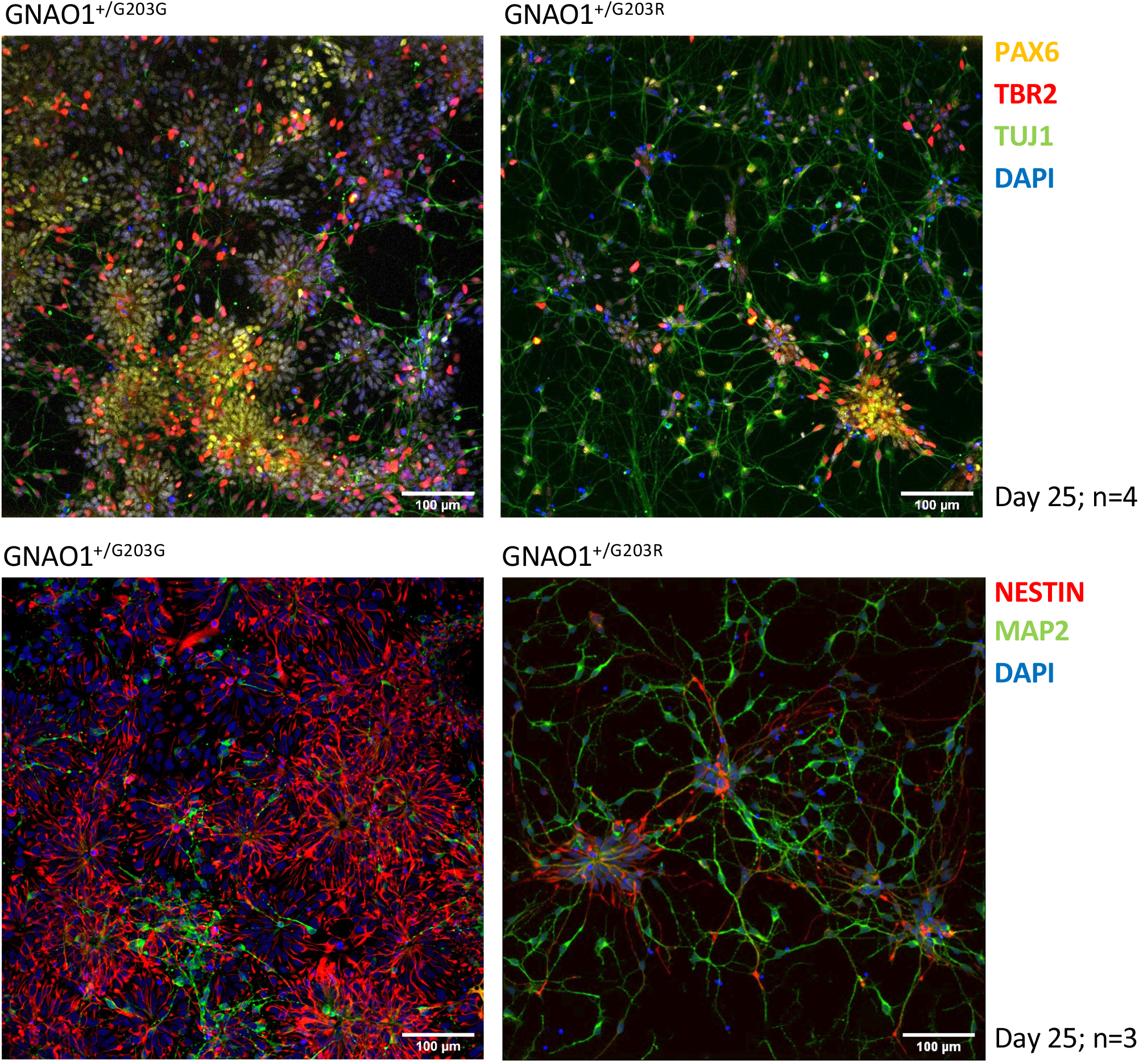
Immunostaining analysis at day 25 of differentiation. Immunostaining analysis with the indicated primary antibodies and DAPI to label nuclei. PAX6 and NESTIN are neural progenitor markers; TBR2 is a neuronal precursor marker; TUJ1 and MAP2 are neuronal markers.

Analysis of the neural rosettes generated from iPSCs collected from a Russian patient with the GNAO1^+/G203R^ variant (herafter GNAO1^+/G203R^#2) showed similar disruption in the morphology of the neural rosettes. GNAO1^+/G203R^#2 hiPSCs, along with the control cell line from a healthy donor, were differentiated into cortical neurons using a method previously described (35). Neural rosette formation was examined on day 30 of differentiation in both control and GNAO1^+/G203R^#2 cultures. The control cultures exhibited cellular rosettes formed by the accumulation of several tens to hundreds of cells, whereas the GNAO1^+/G203R^#2 cultures showed clusters of only 10-20 cells (Figure 4A,B). Immunocytochemical analysis of the neuronal markers TBR1, TUJ1 (Figure 4B), and MAP2 (Figure 4C) revealed the presence of positively stained cells at the edge of the neural rosettes in the control culture. In contrast, the number of these cells was significantly lower in the patient line-derived culture (Figure 4B,C). Similarly, a marked reduction in NESTIN-positive progenitor cells was observed in the variant culture compared with the control (Figure 4C). In addition, the variant-containing rosettes exhibited disrupted cytoarchitecture, a lack of cell polarization, and the absence of radial structure. Apical polarization of β-catenin was also impaired in the variant-containing culture (Figure 4D).

**Figure 4.**
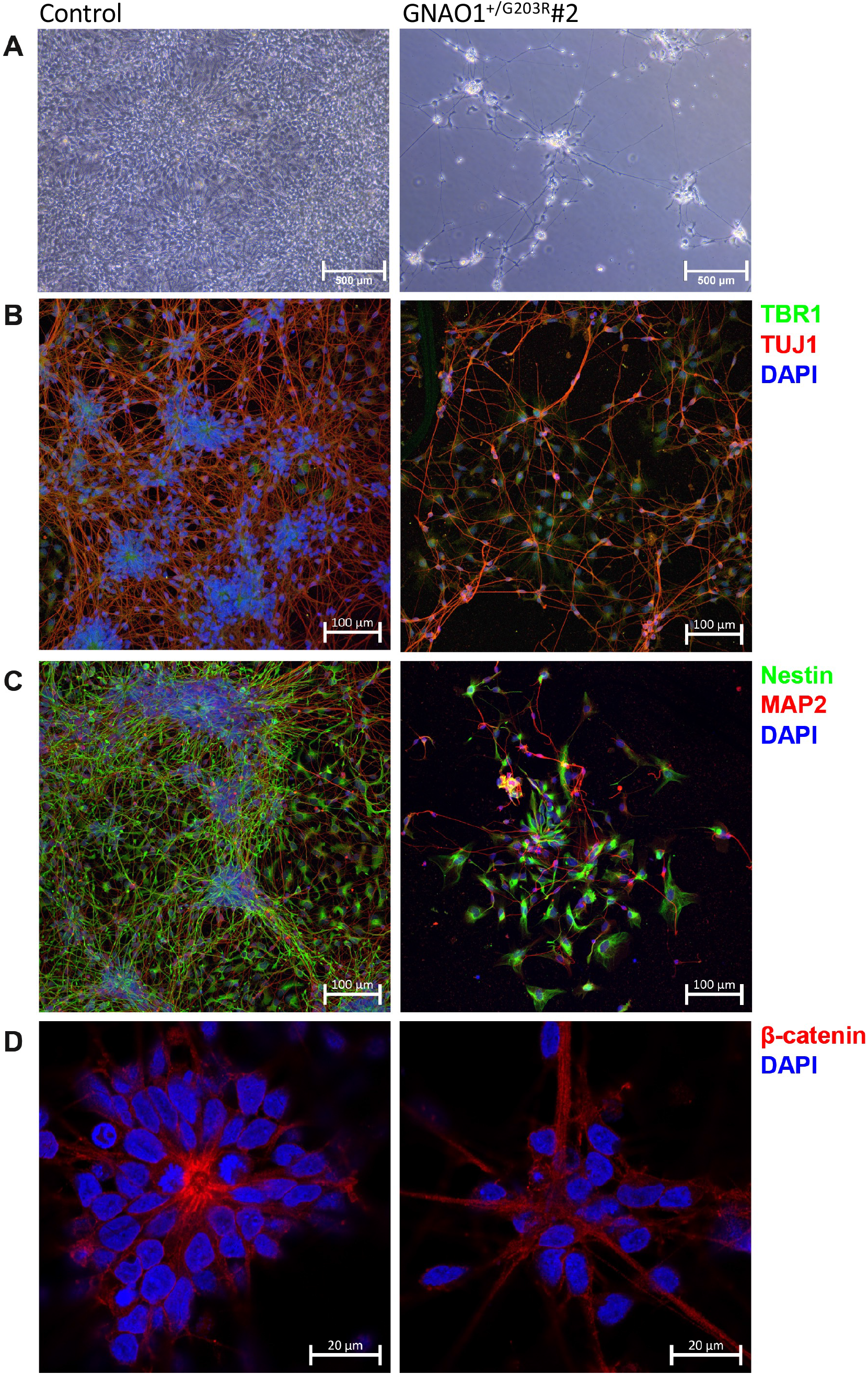
Analysis of neuronal rosettes morphology in the independent GNAO1^+/G203R^#2 line. **A.** Phase contrast images of GNAO1^+/G203R^#2 cells at the neural rosette stage, corresponding to day 30 of differentiation. Note that the in the protocol used for differentiation of GNAO1^+/G203R^#2 cells (as described in the Methods section) has a different timing from the one depicted in figure 2A. **B-D**. Immunostaining was performed using the specified primary antibodies along with DAPI for nuclear labeling. TBR1 is a marker of early-born neurons (B); and TUJ1 (B) and MAP2 (C) are markers for neurons; NESTIN serves as a marker for neural progenitors (C); β-catenin as a marker of apical polarization in neural rosettes (D). Number of individual cultures for experiments was 3.

Collectively, these data suggest an early impairment that affects *GNAO1* p.G203R neural progenitor cells. Importantly, morphologic abnormalities in neural rosettes were consistently observed in two independent patient-derived *GNAO1* p.G203R lines.

### The *GNAO1* p.G203R substitution alters neuronal maturation

At day 40, corresponding to a stage in which differentiated cells have entered the neuronal maturation stage, GNAO1^+/G203R^-derived neurons continued to express lower levels of *TBR1*, which highlights a reduction in the presence of layer VI neurons (Figure 5A,B). On the other hand, they tended to express higher levels of *GFAP*, an astrocyte marker, compared to the isogenic control, and failed to downregulate *PAX6* expression, suggesting an increased amount of astrocyte and progenitor cells (Figure 5A,B). Furthermore, the expression levels of *FOXG1*, a telencephalic gene that plays an important role in neural differentiation and in balancing the excitatory/inhibitory network activity, were increased (Figure 5A). At later time points (day 50 and day 70), we confirmed significant *TBR1* downregulation in variant-containing neural cultures, whereas we observed an increased trend of upregulation for *GFAP* (at both day 50 and day 70), *PAX6* (day 50, not tested at day 70) and *FOXG1* (only at day 50), despite statistical significance not always being reached (Figure 6A-C).

**Figure 5.**
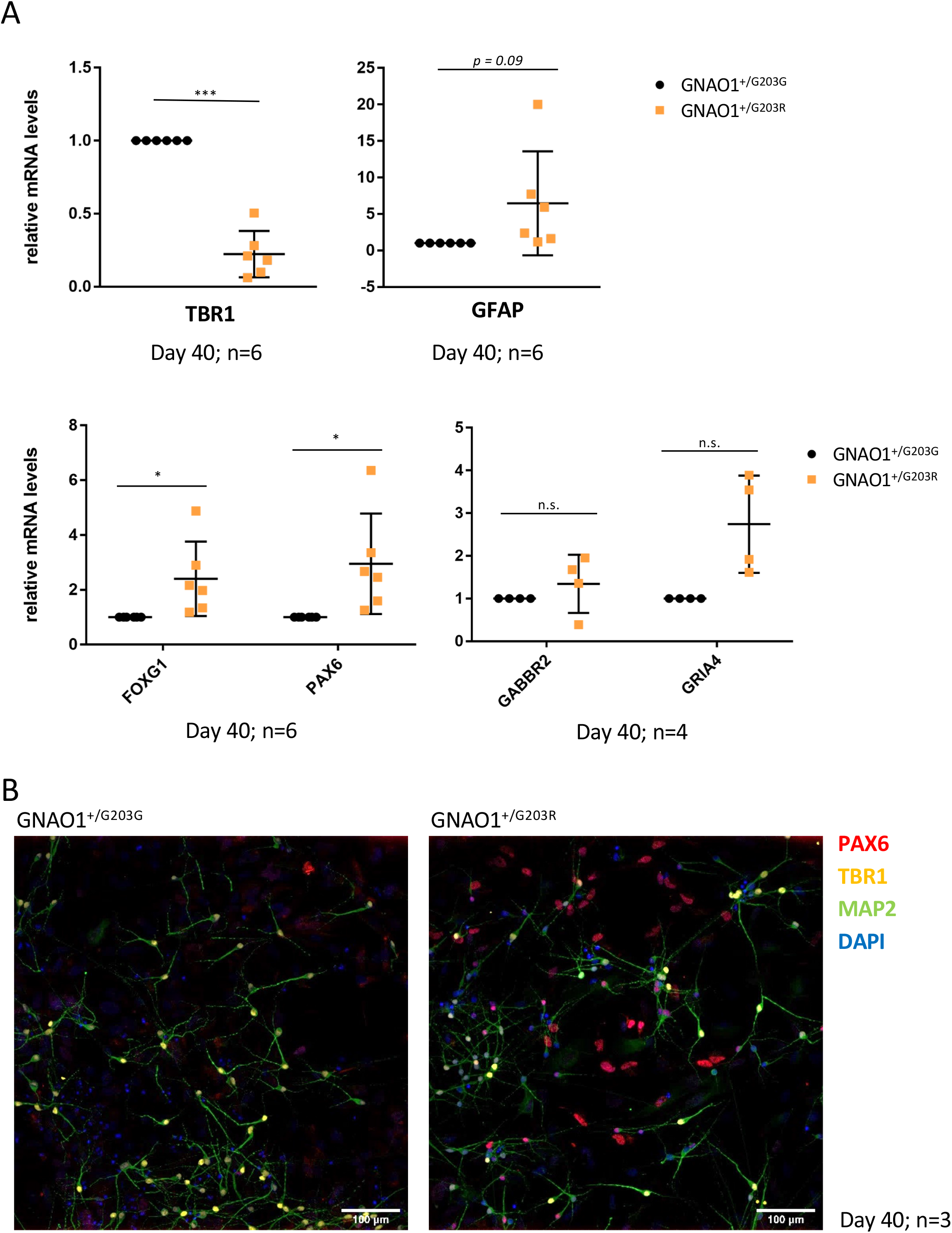
Marker expression analysis at day 40 of differentiation. **A.** Real-time qRT-PCR analysis of the expression of the indicated markers in differentiating cells. The graphs show the average and standard deviation (Student’s t test; paired; two tails; *p<0.05, ***p<0.001, n.s. nonsignificant). **B.** Immunostaining analysis with the indicated primary antibodies and DAPI to label nuclei.

**Figure 6.**
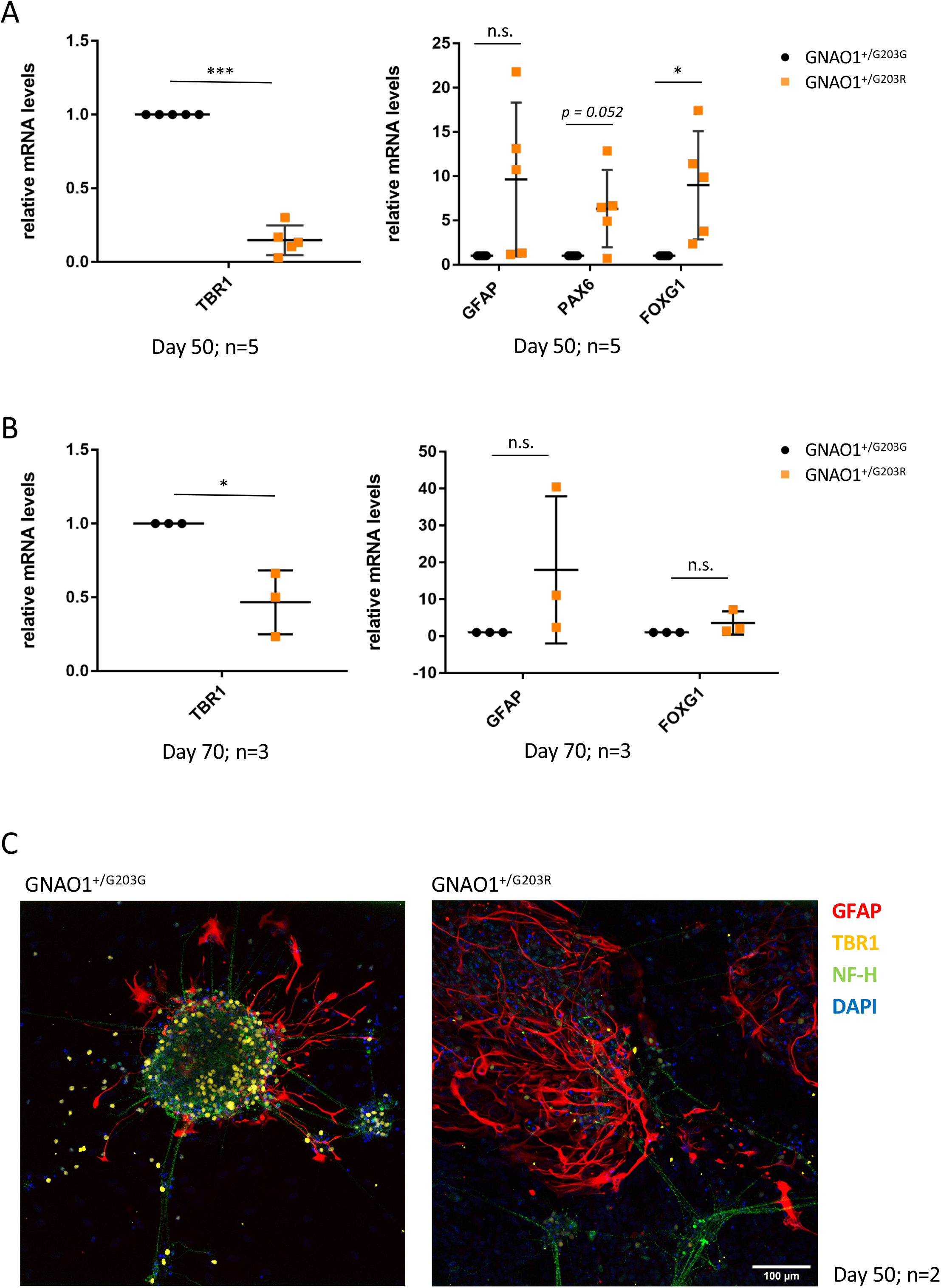
Marker expression analysis at day 50 and 70 of differentiation. **A-B.** Real-time qRT-PCR analysis of the expression of the indicated markers in differentiating cells at the indicated time points. The graphs show the average and standard deviation (Student’s t test; paired; two tails; *p<0.05, ***p<0.001, n.s. nonsignificant). **C.** Immunostaining analysis with the indicated primary antibodies and DAPI to label nuclei.

Collectively, these findings indicate that the *GNAO1* p.G203R variant leads to altered gene expression as well as premature and defective differentiation processes affecting both neurons and astrocytes.

### Spontaneous and evoked calcium transients were reduced in *GNAO1* p.G203R cortical neurons

Functional characterization via calcium imaging was performed on days 40 and 50. As shown in Figures 7 and 8, this analysis suggests impaired functional maturation in neuronal cultures obtained from patient-derived GNAO1^+/G203R^ iPSCs. At both time points, GNAO1^+/G203R^-derived neurons showed decreased levels of basal intracellular free calcium concentration ([Ca^2+^]_i_) and a reduced fraction of neurons with spontaneous activity (Figure 7B). In line with this finding, calcium-dependent signaling was affected in GNAO1^+/G203R^ neurons at day 50, which displayed lower levels of pGSK3β at the inhibitory Ser9 residue compared to control neurons (Supplementary Figure S3). We then tested the response of control and variant-containing neurons to different neurotransmitters or selective agonists. The highest response levels were obtained, both in control and GNAO1^+/G203R^-derived neurons, upon application of glutamate (Figure 8A), but with a significant decrease in the percentage (at day 40) and in the amplitude of the responses in GNAO1^+/G203R^-derived neurons (at both time points). The most striking difference was detected upon stimulation with GABA (Figure 8B). While the vast majority of wild-type neurons responded to GABA stimulation at both time points, this effect was almost completely abolished by the pathogenic variant. Moreover, the amplitude of the [Ca^2+^]_i_ increase was significantly lower even in the few GNAO1^+/G203R^ cells that were able to respond to GABA. We also tested baclofen, a specific agonist of GABA_B_ receptors (Figure 8C). In this case, we observed a statistically significant difference only in the number of responding cells at day 50. Reduced activity did not correlate with a decrease in *GABBR2* (GABA_B_ Receptor 2) and *GRIA4* (Glutamate Ionotropic Receptor AMPA Type Subunit 2) mRNA levels (Figure 5A), pointing to functional impairment rather than decreased expression of receptors. Reduced activity in response to other neurotransmitters, such as glycine, acetylcholine, and ATP was also observed in terms of the number of responding cells (Figure 8D-F). These results suggest that the *GNAO1* p.G203R variant leads to a delay in neuronal maturation, which in turn causes a profound impairment of neuronal function.

**Figure 7.**
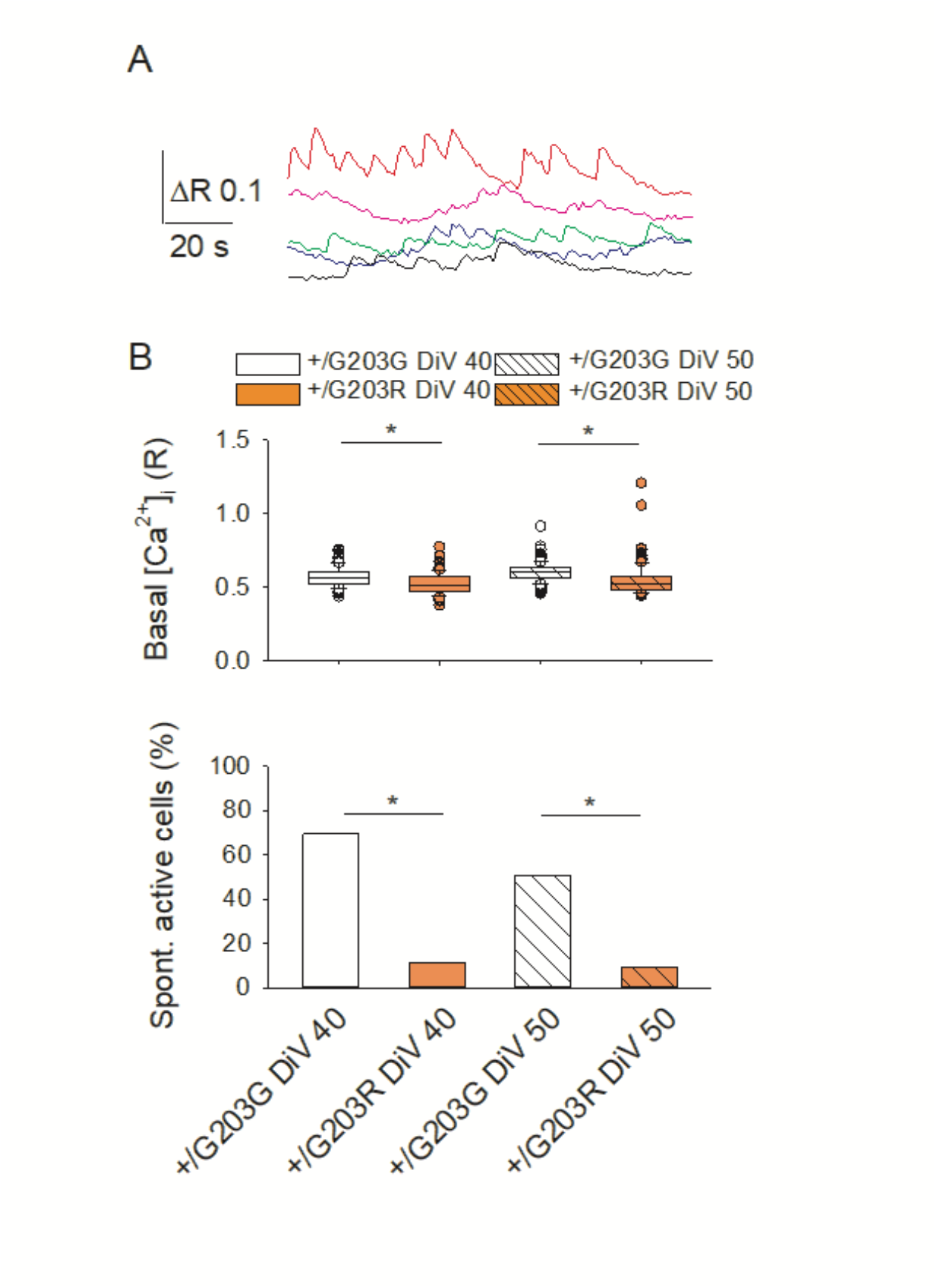
Neurons carrying the p.G203R GNAO1 substitution exhibit lower values of basal [Ca^2+^]_i_ and a reduced spontaneous Ca^2+^ activity. **A.** representative spontaneous Ca^2+^ transients recorded in control cells. **B.** up, [Ca^2+^]_i_ basal values measured in individual cells. Please note the reduced basal [Ca^2+^]_i_ for p.G203R mutant neurons. Number of examined cells: 160, 159, 229 and 230, for p.G203G DIV40, p.G203R DIV40, p.G203G DIV50 and p.G203R DIV50, respectively. Bottom, percentage of neurons exhibiting spontaneous Ca^2+^ transients, same cells as upper panel. *, p<0.001.

**Figure 8.**
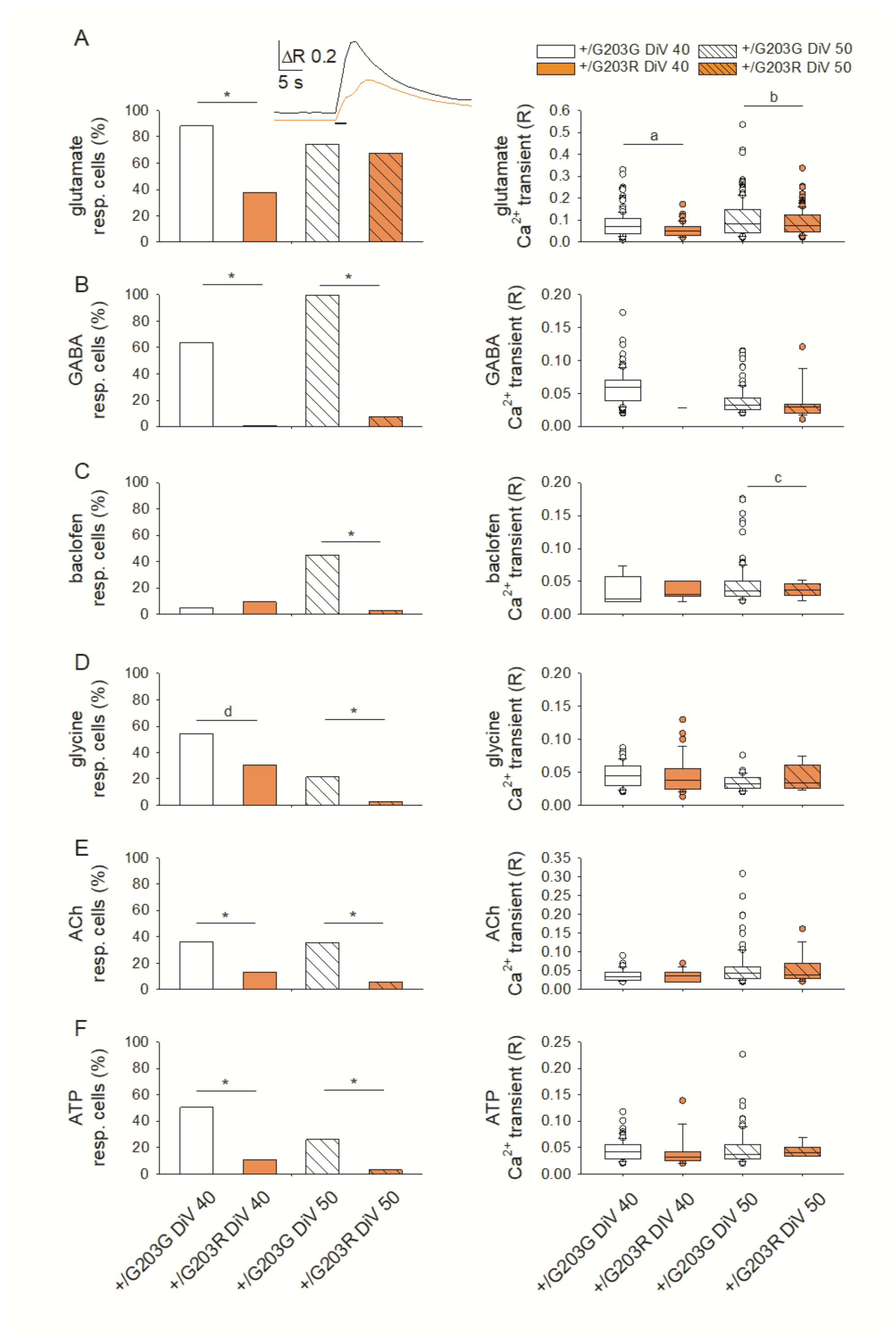
The p.G203R GNAO1 substitution reduces the response of differentiated neurons to several neurotransmitters in terms of [Ca^2+^]_i_ elevations. **A.** left, percentage of neurons exhibiting Ca^2+^ transients induced by the application of glutamate (1 mM, 3 s); inset, representative Ca^2+^ transients elicited by glutamate administration (1 mM, 3 s), in a control (black) and GNAO1 p.G203R (red) iPSC-derived neuronal cells. Number of examined cells: 160, 159, 229 and 230, for p.G203G DIV40, p.G203R DIV40, p.G203G DIV50 and p.G203R DIV50, respectively. Right, amplitude of the [Ca^2+^]_i_ increase induced by glutamate application in the responding cells. **B.** left, percentage of neurons exhibiting Ca^2+^ transients induced by the application of GABA (100 μM, 3 s). Same cells as fig. 7A. Right, amplitude of the [Ca^2+^]_i_ increase induced by GABA application in the responding cells. **C.** left, percentage of neurons exhibiting Ca^2+^ transients induced by the application of baclofen (100 μM, 3 s). Same cells as fig. 7A. Right, amplitude of the [Ca^2+^]_i_ increase induced by baclofen application in the responding cells. **D.** left, percentage of neurons exhibiting Ca^2+^ transients induced by the application of glycine (30 μM, 3 s). Same cells as fig. 7A. Right, amplitude of the [Ca^2+^]_i_ increase induced by glycine application in the responding cells. **E.** left, percentage of neurons exhibiting Ca^2+^ transients induced by the application of ACh (100 μM, 3 s). Same cells as fig. 7A. Right, amplitude of the [Ca^2+^]_i_ increase induced by ACh application in the responding cells. **F**. left, percentage of neurons exhibiting Ca^2+^ transients induced by the application of ATP (100 μM, 3 s). Same cells as fig. 7A. Right, amplitude of the [Ca^2+^]_i_ increase induced by ATP application in the responding cells. *, p<0.001; a, p=0.008; b, p=0.022; c, p=0.006, d, p=0.007.

## DISCUSSION

In this study, we provide a side-by-side comparison of patient-derived hiPSCs and an isogenic control in which the p.G203R change has been reverted to WT (GNAO1^+/G203G^). The highly conserved glycine 203 residue represents a pathogenic variant hotspot, associated with a mixed, severe phenotype characterized by epileptic seizures and movement disorders (36;37). The major clinical signs associated with the *GNAO1* p.G203R variant are neonatal/infantile-onset therapy-resistant epilepsy (generalized or focal), axial hypotonia, dystonia, and hyperkinetic movement disorders (chorea associated with paroxysmal events requiring intensive care). Together with glutamic acid 246 and arginine 209, this residue plays a crucial role in the allosteric regulation mediating Gβγ release (38). The core molecular dysfunction for the three most common *GNAO1* pathogenic variants (p.G203R, p.R209C/H, and p.E246K) is the constitutive GTP-bound state of the mutant Gαo proteins coupled to their inability to adopt the active conformation which allow proper binding to regulators and effectors (24). Indeed, these three residues are located in the switch II or switch III regions, which are important for guanidine nucleotide-dependent regulation of downstream effectors.

In this work, the GNAO1^+/G203R^ hiPSC line from an American patient and the GNAO1^+/G203G^ isogenic control were differentiated into cortical neurons using a method that recapitulates human corticogenesis. The comparison with the isogenic control shows that p.G203R is sufficient to impair the timeline of differentiation as early as at day 25, representing the onset of neuronal differentiation from neural progenitor cells. Similar impairment in the ability to form neural rosettes was consistently observed in a second GNAO1^+/G203R^ line derived from an unrelated Russian patient. This agreement confirms the importance of the hiPSCs *GNAO1* model in elucidating the pathogenesis of the disease and underscores the importance of further investigation of the role of this variant in the pathogenesis of *GNAO1*-related disorders. This neurodevelopmental phenotype is clinically relevant because, based on imaging, microcephaly, progressive and diffuse cerebral/cerebellar atrophy, hypomyelination/delayed myelination, and abnormalities of the basal ganglia (e.g., hypoplasia/atrophy of the caudate nucleus) or corpus callosum (thin corpus callosum) have been observed in the most severe cases harboring the p.G203R substitution (39).

Altered timing of proliferation and differentiation has been reported in many neurodevelopmental disease models, such as Fragile X syndrome (40;41), Tuberous Sclerosis Complex (42), and CDKL5 Deficiency Disorder (43). The neurodevelopmental phenotype observed here is in line with previous observations from *GNAO1* animal models. We have recently shown that there is a decrease in neural progenitor cells in two different mouse models of *GNAO1*-related disorders (p.C215Y and p.G203R), suggesting an impairment in neuronal differentiation or migration, specifically in the developing cerebral cortex (13). We found enlargement of lateral ventricles and altered motor cortex thickness.

We also reported a reduction of cell number in the total cortex, while no change was observed in the striatum and hippocampus. Accordingly, here we show that compared to the isogenic control line, human *GNAO1* p.G203R differentiating neural progenitors are characterized by reduced expression of early neural genes, such as *NESTIN*, *OTX2* and *PAX6*. Moreover, in line with the mouse model which has half of the number of *Tbr2*-positive cells in the ventricular and subventricular zones and has a lower cell density in layer VI of the cortex (*tbr1*-positive cells) (22), here we found decreased levels of both *TBR1* and *TBR2* transcripts in the iPSC-derived neurons. Altered expression of *FOXG1*, which was significantly upregulated at day 40 and shows the same trend at later time points, is intriguing. Indeed, variants in the *FOXG1* gene have been linked to a rare form of epileptic encephalopathy (OMIM #613454) (44). *TBR1* is a master regulator of cortex development that plays a crucial role in the formation of neocortical layer VI, regulating both neuronal migration and synaptic development (45;46). Pathogenic variants in *TBR1* gene are associated with cortical malformations, intellectual disability and autism spectrum disorder. Furthermore, it has been shown that *TBR1* regulates neural stem cell fate by inhibiting astrocytes formation (47). Interestingly, we report increased expression of *GFAP*, an intermediate progenitor marker expressed in astrocytes, suggesting that *GNAO1* pathogenic variants could have an effect both on neuronal and glial populations. *GFAP* overexpression has been found both in neurodevelopmental and neurodegenerative diseases, such as Fragile X syndrome and Alzheimer’s disease (33;48;49;50;51). Moreover, abnormal astrocytes proliferation linked to altered timing of neurogenesis and astrogenesis has been observed in RASopathies and Down Syndrome, further suggesting the role of astrocytes on the onset of neurodevelopmental diseases (52). Increased GFAP expression is also considered as a hallmark of reactive astrocytes, which are astrocytes that underwent morphological and functional modification as a consequence of brain injury or disease (53). Notably, since prolonged astrogliosis contributes to the epileptic phenotype (54) and astrocytes alteration has been observed in different DEE models (55;56;57), astrocytes are considered a target for therapeutic approaches in epilepsy (58). Our present findings suggest that this might be extended to *GNAO1*-related disorders as well.

Our functional analysis supports the concept that the *GNAO1* p.G203R substitution leads to a lower degree of neuronal maturation. Control neurons exhibit sustained spontaneous [Ca^2+^]_i_ activity, which is significantly reduced in GNAO1^+/G203R^-derived cells, in association with a reduction of the basal [Ca^2+^]_I_ level. As a probable consequence of the decrease in [Ca^2+^]_i_, we found lower levels of pGSK3β(S9) in day 50 differentiated p.G203R neurons compared to control cells, which is in line with previous findings in neuronal cultures with knockout for the *GNAO1* gene or carrying the *GNAO1* p.G203R variant (25). Considering the key role of Wnt-GSK3 signaling in the regulation of axon extension and branching (59), further studies are needed to explore the impact of aberrant GSK3β phosphorylation in p.G203R neuronal morphogenesis. Furthermore, GNAO1^+/G203R^-derived neurons are less responsive to several agonists, indicating a lower functional level of different crucial receptor systems. In particular, both GABA and glutamate induce a much lower response, suggesting a profound impairment of the neuronal function in Day 40 and Day 50 differentiated p.G203R neurons compared to control cells. Reduced sensitivity to neurotransmitters, and in particular GABA, is observed in several epileptic disorders (60). The functional impairment observed in p.G203R neurons (loss of spontaneous Ca^2+^ transient activity, lower responsiveness to most neurotransmitters) strongly suggests a slower maturation of the molecular machinery necessary for proper synaptic connections, likely leading to dysfunctional cortical and subcortical circuitry, and consequently to the typical clinical signs, such as drug-resistant neonatal epilepsy and hyperkinetic movement disorders.

The hiPSC-based model described here will be used as an *in vitro* platform for translational studies. Both the neurodevelopmental phenotype and the functional impairment represent possible readouts to evaluate the effects of promising new drug candidates, such as caffeine and zinc (19;24), and new gene therapy approaches, such as those based on recombinant adeno-associated virus (rAAV)-mediated delivery of genetic constructs encoding WT *GNAO1*, as well as short hairpin RNA (shRNA) to suppress the endogenous gene (61). A major limitation of the present work, however, lies in the absence of additional patient-derived hiPSC lines. Future investigations will be imperative to determine whether the phenotypes observed in p.G203R cells are replicated in the presence of different *GNAO1* pathogenic variants.

In summary, our study solidifies the understanding of GNAO1 encephalopathy as a neurodevelopmental disorder, shedding light on the specific molecular and functional disruptions caused by the p.G203R variant. Our findings illuminate critical aspects of impaired neurogenesis, altered neuronal differentiation, and dysfunctional responses in affected neural cells. While this research delves into the early stages of neurodevelopment, further investigations into the variant’s impact on fully matured neurons are imperative. Additionally, exploring a broader spectrum of *GNAO1* pathogenic variants associated with diverse clinical features will enhance our comprehension of this complex disorder. These avenues of study not only promise to deepen our understanding of *GNAO1*-related neurological conditions but also pave the way for targeted interventions and therapeutic strategies.

## MATERIALS AND METHODS

### Reprogramming of patient-derived hiPSCs and generation of the gene edited GNAO1^+/G203G^ control line

GNAO1^+/G203R^ iPSCs derived from an American patient and the isogenic control line were kindly provided by the nonprofit organization Child’s Cure Genetic Research (Fremont, CA, USA). Briefly, fibroblasts were derived at the Stanford Children’s Hospital (CA, USA) from a skin punch biopsy of a GNAO1^+/G203R^ patient. The patient (male, born May 2017, 2 years old at the time of biopsy) began to exhibit seizures at the age of 6 weeks and epilepsy was controlled with anti-epileptic drugs. Overtime seizures transformed to infantile at the age 4 months and he was treated with adrenocorticotropic hormone (ACTH) and vigabatrin. Seizures have been in remission until the age of 6 years old. Patient exhibited global development delay, weak muscle tone, movement disorders (chorea and dystonia that were more pronounced with fever) and limited communication skills through visual eye gaze device. Reprogramming to iPSCs, followed by standard characterization, was performed by Applied StemCell Inc. The isogenic control line was generated using CRISPR/Cas9-mediated genome editing by the Genome Engineering & Stem Cell Center (GESC@MGI) at Washington University in St. Louis (https://GeneEditing.wustl.edu).

GNAO1^+/G203R^#2 hiPSCs were generated by reprogramming of skin fibroblasts from a patient from Moscow with GNAO1 encephalopathy carrying the c.607 G>A (p.Gly203Arg) variant, as described in (61), which were kindly provided by Dr. Bardina (Institute of Gene Biology, Russian Academy of Sciences, Moscow) upon the proper written patient consent. The patient was born in October 2016 (male, 1 year old at the time of biopsy). From day 7 after birth, epileptic seizures were noted in the form of bilateral tonic seizures. Remission of seizures was achieved at 5 months of age upon treatment with levetiracetam and vigabatrin. In addition to seizures, the patient presented with a complex set of symptoms, including psychomotor developmental delay, choreiform hyperkinesia, and dystonic hyperkinesia. To generate control fibroblasts, a skin biopsy was obtained from a healthy 26-year-old female donor after informed consent. The biopsy was cut into small pieces and cultured under the coverslip in DMEM (PanEco, Russia) with 10% FBS (Hyclone, USA), 2 mM Glutamine (PanEco), 1X penicillin-streptomycin (PanEco). Medium was changed once in three days. After seven days, fibroblasts were passaged with 0.25% Trypsin (Gibco). Human skin fibroblasts from the patient and the healthy donor were reprogrammed using the protocol described in Shuvalolva et al, 2020 (62).

### RNA-Sequencing and bioinformatic analysis

FASTQ files from paired-end RNA-Seq analysis performed using the Illumina® Stranded Total RNA Prep kit with Ribo-Zero Plus treatment on GNAO1^+/G203R^ and GNAO1^+/G203G^ hiPSC-derived NPCs at day 7, generated at Rarebase PBC, and day 11, generated as previously described in Whye et al. (2023) (basic protocol 1 and support protocol 3) (63). For each time point, 3 replicates from one differentiation batch were used. Raw RNA-Seq datasets are available at the NCBI Sequence Read Archive (SRA) with the accession number PRJNA991628. Forward and reverse reads were aligned to a database of rRNA sequences using Bowtie 2 v2.4.4 (64) with --nofw and --nofw parameters, respectively; only read pairs in which both mates did not map to these abundant molecules were retained for downstream analyses. STAR v2.7.10b (65) with default mapping parameters was used to align reads to the human GRCh38 genome, providing the GENCODE 42 (66) GTF file to generate a splice junction database and to calculate gene-level read counts with the -- quantMode TranscriptomeSAM GeneCounts option. Read counts reported in the fourth column of ReadsPerGene.out.tab files were combined into a count matrix using a custom Python script. This matrix was used to calculate normalized expression values, expressed as FPKM (Fragments Per Kilobase of exon per Million mapped reads), using the “fpkm” function from the DESeq2 v1.34.0 R package (67). We retained only protein-coding or lncRNA genes with FPKM > 0.5 in at least three samples for subsequent analysis. Prior to performing PCA using the DESeq2 “plotPCA” function, we applied regularized-log (rlog) transformation to the count data. To identify genes which are differentially express (DEGs) between GNAO1^+/G203R^ and GNAO1^+/G203G^ samples at both time points, Wald test with independent filtering was performed using the “DESeq” function. Apeglm method (68) was employed for log2(fold change) shrinkage. The FDR-adjusted p-value threshold to select DEGs was set to 0.01. Additionally, in both contrasts, we only selected genes that had an average FPKM greater than 0.5 in at least one of the two conditions being compared. The intersections between the DEGs identified in the two comparisons were visualized using the UpSetR v1.4.0 R package (69). Gene expression heatmaps were generated with the ComplexHeatmap v2.10 R package (70) using row-scaled (Z-score), rlog-transformed count data. The WebGestaltR v0.4.4 R package (71) was employed to perform Gene Set Enrichment Analysis (GSEA), using shrunken log2(fold change) values as a ranking metric.

### HiPSC culture and differentiation

GNAO1^+/G203R^ iPSCs were cultured in Nutristem-XF (Biological Industries) supplemented with 0.1% Penicillin-Streptomycin (Thermo Fisher Scientific) onto Biolaminin 521 LN (Biolamina) functionalized plates. The culture medium was refreshed every day and cells were passaged every 4–5 days using 1 mg/mL Dispase II (Thermo Fisher Scientific). Cells were routinely tested for mycoplasma contamination. GNAO1^+/G203R^ hiPSCs were differentiated into cortical neurons as previously described (33).

GNAO1^+/G203R^#2 hiPSCs were cultured in mTeSR™1 medium (Stem Cell Technologies) on Matrigel (Corning)-coated plates following the manufacturer’s guidelines. Every day, the culture medium was renewed, and cells were passaged every 4-5 days up to 70-85% confluency utilizing 0.5 mM EDTA (Thermo Fisher Scientific). On the first day post-subculturing, cells were cultured in medium supplemented with 5 μM of the ROCK inhibitor StemMACS™ Y27632 (Miltenyi Biotec). The differentiation of hiPSCs into midbrain neurons was performed as previously described (35). The differentiation of GNAO1^+/G203R^#2 hiPSCs into mature human neurons comprised three main stages: (1) hiPSC differentiation into neuronal progenitor cells (NPCs); (2) expansion and maturation of NPCs; and (3) differentiation of NPCs into mature neurons. In brief, hiPSCs were differentiated into NPCs using the commercially available PSC Neural Induction Medium (Thermo Fisher Scientific) following the manufacturer’s guidelines. This was considered as day 0 (D0). After 10 days of induction to NPCs, cells were detached from the substrate with 0.5 mM EDTA and were plated at density of 250,000-400,000 cells per cm^2^ in Petri dishes or multi-well plates coated with Matrigel. Further cultivation took place in the medium for expansion and maturation of NPCs consisting of advanced DMEM/F12 (Thermo Fisher Scientific), Neurobasal Medium (Thermo Fisher Scientific), 2% Neural Induction Supplement (GIBCO). The ROCK inhibitor Y27632 5 μM was used for every seeding overnight with the following replace to the medium without ROCK inhibitor. After 18 days the differentiation protocol for maturation into neurons was initiated. Neural precursors were cultured and frozen until the third passage. We reseeded neural precursors to a density of 300,000-400,000 cells per cm^2^. NPCs were seeded into wells coated with Matrigel and cultured in the medium containing DMEM/F12 (Thermo Fisher Scientific), 1X GlutaMAX (Thermo Fisher Scientific), 2% B27 (Thermo Fisher Scientific), 1X Penicilin-Streptomicin (Thermo Fisher Scientific), BDNF 20 ng/mL (GenScript), GDNF 20 ng/mL (GenScript), 200 μМ ascorbic acid (Sigma Aldrich), 4 μМ forskolin (Sigma Aldrich) for 2 weeks with subculturing during the first week. The ROCK inhibitor was used every passage overnight with the following replacement to the medium without the ROCK inhibitor.

### Real-time PCR

RNA was extraction and Real-time RT-PCR was performed as described in Brighi et al. 2021 (33). A complete list of primers, including the housekeeping control gene *ATP5O*, is provided in Supplementary material (Supplementary Table S2).

### Immunostaining

GNAO1^+/G203R^ differentiating cells were immunostained as described in Brighi et al. 2021 (33) using the following primary antibodies: mouse anti-PAX6 (sc81649 Santa Cruz Biotechnology, 1:50), mouse anti-GFAP (MAB360 Merck Life Sciences, 1:500), chicken anti-MAP2 (ab5392 Abcam, 1:2000), rabbit anti-β-TUBULIN III (TUJ1) (T2200 Merck Life Sciences, 1:2000), Rabbit Anti-Nestin (MA5-32272 Thermo Fisher Scientific, 1:100), Rabbit Anti-TBR2 (ab15894 Millipore,1:50). AlexaFluor secondary antibodies (Thermo Fisher Scientific) were used at the concentration of 1:250, and DAPI (Merck Life Sciences) was used to stain nuclei. An inverted Olympus iX73 microscope equipped with an X-Light V3 spinning disc head (Crest Optics), a LDI laser illuminator (89 North), using 405nm, 470nm, 555nm and 640nm wavelengths, depending on the fluorophores, a Prime BSI sCMOS camera (Photometrics) and MetaMorph software (Molecular Devices) with a 20x objective (Olympus), was used for image acquisition.

GNAO1^+/G203R^#2 cells at the stage of neuronal rosettes (30 day of differentiation) were fixed in 4% paraformaldehyde solution (Merck Life Sciences) for 15 min at room temperature and washed thrice with 1X PBS (Thermo Fisher Scientific). Fixed cells were permeabilized for 1 min using ice-cold PBS supplemented with 0.1% Triton X-100, blocked for 30 min with PBS supplemented with 1% BSA. Cells were then incubated overnight at 4°C with primary antibodies at the following dilutions: rabbit anti-MAP2 (ab254264 Abcam, 1:2000), mouse anti-βTUBULIN III (TUJ1) (T8660 Sigma-Aldrich, 1:500), mouse Anti-Nestin (ab6142 Abcam, 1:250), rabbit anti-TBR1 (ab183032 Abcam,1:200), mouse anti-β-Catenin (571-781 BD Transduction Laboratories, 1:200). The day after, the primary antibody solution was washed out and the cells were incubated with secondary antibodies (Alexa Fluor 488 or Alexa Fluor 594 conjugates, Jackson ImmunoResearch, 1:250) for 2 h at room temperature in the dark. DAPI (Merck Life Sciences) was used for nucleus staining. Images were acquired with an LSM710 confocal microscope.

### Western Blotting

Differentiating cells were lysed with RIPA Buffer with freshly added protease and phosphatase inhibitors (Thermo Fisher Scientific), and kept on ice for 30 min. Afterwards they were centrifuged at 12.000×g (20 min, 4 °C), the pellet was discarded and the supernatant transferred into new tubes, and stored at -80 °C. Protein content was determined using the BCA protein Assay (Thermo Fisher Scientific). Protein extracts (20 μg) were resuspended in Novex Tris-Glycine SDS Sample Buffer (Thermo Fisher Scientific) containing Sample Reducing Agent, heated for 5 min at 85 °C, and separated on precast Novex Tris-Glycine Mini Gels. Proteins were transferred using iBlot™ 2 on a PVDF membrane, which was then blocked in 5% nonfat dry milk (Biorad Laboratories) in T-TBS for 1h at RT. Membranes were then incubated O/N with anti-Phospho-Gsk3β (Ser9 - D85E12) rabbit monoclonal antibody (1:1.000, cat#5558, Cell Signaling), anti-GSK-3β (3D10) mouse monoclonal antibody (1:1.000, cat#9832, Cell Signaling), and anti-GAPDH (6C5) mouse monoclonal antibody (1:500, cat# sc-32233, Santa Cruz Biotechnologies) in T-TBS with 5% BSA. After incubation, membranes were washed three times in T-TBS for 10 min at RT, and incubated for 1h with anti-mouse horseradish peroxidase-conjugated antibodies (1:10.000, cat#115-035-003, Jackson Immunoresearch Laboratories), or anti-rabbit horseradish peroxidase-conjugated antibodies (1:10.000, cat#111-035-003, Jackson Immunoresearch Laboratories) at RT. Reactivity was detected using Western Bright ECL spray device (cat#K-12049-D50, Advansta, Aurogene) and Fluorchem E (ProteinSimple). Densitometric analysis was performed using the ImageJ software (http://rsb.info.nih.gov/ij/).

### Calcium imaging

Changes in free intracellular Ca^2+^ concentration [Ca^2+^]_i_ were quantified by time-resolved digital fluorescence microscopy using the Ca^2+^ indicator Fura-2 (excitation 340 nm and 380 nm, emission 510 nm). The changes of [Ca^2+^]_i_ were expressed as R = F340/F380. iPSCs-derived neuronal cells were incubated with the cell-permeant Fura-2 acetoxylmethylester (2 μM; Molecular probes, Life technologies) for 1 h at 37 °C in culture medium. Ca^2+^ responses were elicited by administering the neurotransmitter or the agonist dissolved in normal external solution for 3 s. Cells were continuously perfused during the experiment. [Ca^2+^]_i_ variations were obtained separately from each individual cell, using the MetaFluor 7.0 software (Molecular Devices, USA).

Amplitude values were expressed as means ± S.D. and analyzed using one-way ANOVA test. When necessary, the non-parametric Dunn’s one-way ANOVA on ranks was used. In case of significance, all pairwise multiple comparison procedure was used (Holm-Sidak, or Dunn’s method for non-parametric tests). Responsive cells data were expressed as % and analyzed using chi-square test. The minimum power of statistical tests was set at 0.8. The significance for all tests was set at p < 0.05.

## AUTHOR CONTRIBUTION

Conceptualization, M.C.B., M.A.L., V.L.K., S.M., S.F., A.R.; Formal analysis, M.C.B., T.D., A.C., D.S.; Investigation, M.C.B., T.D., D.S.; Methodology, M.C.B., T.D., A.C., M.M., V.d.T., R.R.N., M.R, Z.B., V.A.B., E.A.V., L.B., A.N.B., R.P., D.W.; Resources: I.M.; Project administration, A.R.; Supervision, E.D.B., G.G.T., M.A.L., V.L.K., S.M., S.F., A.R.; Writing—original draft, A.R.; Writing - Review & Editing, M.C.B., T.D., A.C., E.D.B., V.L.K, S.M., S.F., A.R.. All authors read and approved the final manuscript.

## ACKNOWLEDGEMENTS

The authors wish to thank the Imaging Facility at Center for Life Nano Science, Fondazione Istituto Italiano di Tecnologia, for support and technical advice. We thank the Genome Engineering & Stem Cell Center (GESC@MGI) at the Washington University in St. Louis for cell line engineering services. We thank Federico Salaris and the other members of the Rosa lab for helpful discussion. We thank Koval Alexey and the other members of the Katanaev lab for support and helpful discussions. We thank Dr. Maryana Bardina for providing patient’s fibroblasts. This work was partially supported by “APS Famiglie GNAO1”, Sapienza University of Rome, Fondazione Istituto Italiano di Tecnologia to A.R., by the Russian Science Foundation Grant #21-15-00138 to D.S. and V.L.K., by the Ministry of Science and Higher Education of the Russian Federation grant #075-15-2019-1669 to M.A.L., E.A.V., A.N.B, L.B., and by Istituto Superiore di Sanità (Ricerca Indipendente 2020-2022_ISS20-0ab01a06bd2a) and ISS - Collezione Nazionale di Composti Chimici e Centro di screening (CNCCS) (Rare, Neglected and Poverty Related Diseases – GNAO1 Project) to S.M.

## DATA AVAILABILITY

The RNA-Seq data from GNAO1^+/G203G^ and GNAO1^+/G203R^ NPCs at day 7 and day 11 are available at the NCBI Sequence Read Archive (SRA) with the accession number PRJNA991628.

## REFERENCES

1. Nakamura, K., Kodera, H., Akita, T., Shiina, M., Kato, M., Hoshino, H., Terashima, H., Osaka, H., Nakamura, S., Tohyama, J. and Kumada, T., 2013. De novo mutations in GNAO1, encoding a Gαo subunit of heterotrimeric G proteins, cause epileptic encephalopathy. The American Journal of Human Genetics, 93(3), pp.496–505. doi: 10.1016/j.ajhg.2013.07.014.

2. Ananth, A.L., Robichaux-Viehoever, A., Kim, Y.M., Hanson-Kahn, A., Cox, R., Enns, G.M., Strober, J., Willing, M., Schlaggar, B.L., Wu, Y.W. and Bernstein, J.A., 2016. Clinical course of six children with GNAO1 mutations causing a severe and distinctive movement disorder. Pediatric neurology, 59, pp.81–84. doi: 10.1016/j.pediatrneurol.2016.02.018.

3. Wirth, T., Garone, G., Kurian, M.A., Piton, A., Millan, F., Telegrafi, A., Drouot, N., Rudolf, G., Chelly, J., Marks, W. and Burglen, L., 2022. Highlighting the dystonic phenotype related to GNAO1. Movement Disorders, 37(7), pp.1547–1554. doi: 10.1002/mds.29074.

4. Krenn, M., Sommer, R., Sycha, T. and Zech, M., 2022. GNAO1 haploinsufficiency associated with a mild delayed-onset dystonia phenotype. Movement Disorders, 37(12), pp.2464–2466. doi: 10.1002/mds.29258.

5. Galosi, S., Novelli, M., Di Rocco, M., Flex, E., Messina, E., Pollini, L., Parrini, E., Pisani, F., Guerrini, R., Leuzzi, V. and Martinelli, S., 2023. GNAO1 Haploinsufficiency: The Milder End of the GNAO1 Phenotypic Spectrum. Movement disorders: official journal of the Movement Disorder Society. doi: 10.1002/mds.29585

6. Domínguez-Carral, J., Ludlam, W.G., Junyent Segarra, M., Fornaguera Marti, M., Balsells, S., Muchart, J., Čokolić Petrović, D., Espinoza, I., Ortigoza-Escobar, J.D., Martemyanov, K.A. and GNAO1-Study Group, 2023. Severity of GNAO1-Related Disorder Correlates with Changes in G-Protein Function. Annals of Neurology. doi: 10.1002/ana.26758.

7. Jiang, M., Gold, M.S., Boulay, G., Spicher, K., Peyton, M., Brabet, P., Srinivasan, Y., Rudolph, U., Ellison, G. and Birnbaumer, L., 1998. Multiple neurological abnormalities in mice deficient in the G protein Go. Proceedings of the National Academy of Sciences, 95(6), pp.3269–3274. doi:10.1073/pnas.95.6.3269

8. Wettschureck, N. and Offermanns, S., 2005. Mammalian G proteins and their cell type specific functions. Physiological reviews, 85(4), pp.1159–1204.doi: 10.1152/physrev.00003.2005.

9. Muntean, B.S., Masuho, I., Dao, M., Sutton, L.P., Zucca, S., Iwamoto, H., Patil, D.N., Wang, D., Birnbaumer, L., Blakely, R.D. and Grill, B., 2021. Gαo is a major determinant of cAMP signaling in the pathophysiology of movement disorders. Cell reports, 34(5). doi: 10.1016/j.celrep.2021.108718.

10. Colecraft, H.M., Brody, D.L. and Yue, D.T., 2001. G-protein inhibition of N-and P/Q-type calcium channels: distinctive elementary mechanisms and their functional impact. Journal of Neuroscience, 21(4), pp.1137–1147.doi: 10.1523/JNEUROSCI.21-04-01137.2001.

11. McDavid, S. and Currie, K.P., 2006. G-proteins modulate cumulative inactivation of N-type (Cav2. 2) calcium channels. Journal of Neuroscience, 26(51), pp.13373–13383.doi: 10.1523/JNEUROSCI.3332-06.2006.

12. Zamponi, G.W. and Currie, K.P., 2013. Regulation of CaV2 calcium channels by G protein coupled receptors. Biochimica et Biophysica Acta (BBA)-Biomembranes, 1828(7), pp.1629–1643. doi: 10.1016/j.bbamem.2012.10.004.

13. Worley, P. F., Baraban, J. M., & Snyder, S. H. (1986). Go, a guanine nucleotide-binding protein: immunohistochemical localization in rat brain resembles distribution of second messenger systems. Proceedings of the National Academy of Sciences, 83(12), 4561–4565. doi: 10.1073/pnas.83.12.4561.

14. Guillén, A., Sémériva, M., Bockaert, J. and Homburger, V., 1991. The transduction signalling protein G0 during embryonic development of Drosophila melanogaster. Cellular signalling, 3(4), pp.341–352. doi: 10.1016/0898-6568(91)90063-z.

15. Wolfgang, W. J., Quan, F., Thambi, N., & Forte, M. 1991. Restricted spatial and temporal expression of G-protein α subunits during Drosophila embryogenesis. Development, 113(2), 527–538. doi: 10.1242/dev.113.2.527.

16. Choi, J.M., Kim, S.S., Choi, C.I., Cha, H.L., Oh, H.H., Ghil, S., Lee, Y.D., Birnbaumer, L. and Suh-Kim, H., 2016. Development of the main olfactory system and main olfactory epithelium-dependent male mating behavior are altered in Go-deficient mice. Proceedings of the National Academy of Sciences, 113(39), pp.10974–10979.. 10.1073/pnas.161302611

17. Fremion, F., Astier, M., Zaffran, S., Guillen, A., Homburger, V., & Semeriva, M. 1999. The heterotrimeric protein Go is required for the formation of heart epithelium in Drosophila. The Journal of cell biology, 145(5), 1063–1076. doi: 10.1083/jcb.145.5.1063.

18. Wang, D., Dao, M., Muntean, B.S., Giles, A.C., Martemyanov, K.A. and Grill, B., 2022. Genetic modeling of GNAO1 disorder delineates mechanisms of Gαo dysfunction. Human Molecular Genetics, 31(4), pp.510–522.doi: 10.1093/hmg/ddab235.

19. Di Rocco, M., Galosi, S., Lanza, E., Tosato, F., Caprini, D., Folli, V., Friedman, J., Bocchinfuso, G., Martire, A., Di Schiavi, E. and Leuzzi, V., 2022. Caenorhabditis elegans provides an efficient drug screening platform for GNAO1-related disorders and highlights the potential role of caffeine in controlling dyskinesia. Human Molecular Genetics, 31(6), pp.929–941.doi: 10.1093/hmg/ddab296.

20. Di Rocco, M., Galosi, S., Follo, F.C., Lanza, E., Folli, V., Martire, A., Leuzzi, V. and Martinelli, S., 2023. Phenotypic Assessment of Pathogenic Variants in GNAO1 and Response to Caffeine in C. elegans Models of the Disease. Genes, 14(2), p.319.doi:10.3390/genes14020319.

21. Larrivee, C.L., Feng, H., Quinn, J.A., Shaw, V.S., Leipprandt, J.R., Demireva, E.Y., Xie, H. and Neubig, R.R., 2020. Mice with GNAO1 R209H movement disorder variant display hyperlocomotion alleviated by risperidone. Journal of Pharmacology and Experimental Therapeutics, 373(1), pp.24–33.doi: 10.1124/jpet.119.262733.

22. Silachev, D., Koval, A., Savitsky, M., Padmasola, G., Quairiaux, C., Thorel, F. and Katanaev, V.L., 2022. Mouse models characterize GNAO1 encephalopathy as a neurodevelopmental disorder leading to motor anomalies: from a severe G203R to a milder C215Y mutation. Acta Neuropathologica Communications, 10(1), pp.1–17. doi: 10.1186/s40478-022-01312-z.

23. Savitsky, M., Solis, G.P., Kryuchkov, M. and Katanaev, V.L., 2020. Humanization of Drosophila Gαo to model GNAO1 paediatric encephalopathies. Biomedicines, 8(10), p.395.doi: 10.3390/biomedicines8100395.

24. Larasati, Y.A., Savitsky, M., Koval, A., Solis, G.P., Valnohova, J. and Katanaev, V.L., 2022. Restoration of the GTPase activity and cellular interactions of Gαo mutants by Zn2+ in GNAO1 encephalopathy models. Science Advances, 8(40), p.eabn9350.doi: 10.1126/sciadv.abn9350.

25. Akamine S, Okuzono S, Yamamoto H, Setoyama D, Sagata N, Ohgidani M, Kato TA, Ishitani T, Kato H, Masuda K, Matsushita Y, Ono H, Ishizaki Y, Sanefuji M, Saitsu H, Matsumoto N, Kang D, Kanba S, Nakabeppu Y, Sakai Y, Ohga S. 2020. GNAO1 organizes the cytoskeletal remodeling and firing of developing neurons. The FASEB Journal, 34(12), 16601–16621. doi: 10.1096/fj.202001113R

26. Soldner, F., Laganière, J., Cheng, A. W., Hockemeyer, D., Gao, Q., Alagappan, R., Khurana, V., Golbe, L. I., Myers, R. H., Lindquist, S., et al. 2011. Generation of isogenic pluripotent stem cells differing exclusively at two early onset Parkinson point mutations. Cell, 146, 1358–1367. doi: 10.1016/j.cell.2011.06.019.

27. Sandoe, J. and Eggan, K., 2013. Opportunities and challenges of pluripotent stem cell neurodegenerative disease models. Nature neuroscience, 16(7), pp.780–789. doi: 10.1038/nn.3425

28. Vasques, J.F., Mendez-Otero, R. and Gubert, F., 2020. Modeling ALS using iPSCs: is it possible to reproduce the phenotypic variations observed in patients in vitro?. Regenerative Medicine, 15(7), pp.1919–1933.10.2217/rme-2020-0067

29. Jassal, B., Matthews, L., Viteri, G., Gong, C., Lorente, P., Fabregat, A., Sidiropoulos, K., Cook, J., Gillespie, M., Haw, R. and Loney, F., 2020. The reactome pathway knowledgebase. Nucleic acids research, 48(D1), pp.D498–D503.doi: 10.1093/nar/gkz1031.

30. Bem, J., Brożko, N., Chakraborty, C., Lipiec, M.A., Koziński, K., Nagalski, A., Szewczyk, Ł.M. and Wiśniewska, M.B., 2019. Wnt/β-catenin signaling in brain development and mental disorders: keeping TCF7L2 in mind. FEBS letters, 593(13), pp.1654–1674.doi: 10.1002/1873-3468.13502.

31. Katanaev, V.L., Ponzielli, R., Sémériva, M. and Tomlinson, A., 2005. Trimeric G protein-dependent frizzled signaling in Drosophila. Cell, 120(1), pp.111–122. doi: 10.1016/j.cell.2004.11.014.

32. Koval, A. and Katanaev, V.L., 2011. Wnt3a stimulation elicits G-protein-coupled receptor properties of mammalian Frizzled proteins. Biochemical Journal, 433(3), pp.435–440. doi: 10.1042/BJ20101878.

33. Brighi, C., Salaris, F., Soloperto, A., Cordella, F., Ghirga, S., de Turris, V., Rosito, M., Porceddu, P.F., D’Antoni, C., Reggiani, A. and Rosa, A., 2021. Novel fragile X syndrome 2D and 3D brain models based on human isogenic FMRP-KO iPSCs. Cell Death & Disease, 12(5), p.498. doi: 10.1038/s41419-021-03776-8.

34. Englund, C., Fink, A., Lau, C., Pham, D., Daza, R. A., Bulfone, A., Kowalczyk, T., & Hevner, R. F. (2005). Pax6, Tbr2, and Tbr1 are expressed sequentially by radial glia, intermediate progenitor cells, and postmitotic neurons in developing neocortex. Journal of Neuroscience, 25(1), 247–251. doi: 10.1523/JNEUROSCI.2899-04.2005.

35. Vigont, V.A., Grekhnev, D.A., Lebedeva, O.S., Gusev, K.O., Volovikov, E.A., Skopin, A.Y., Bogomazova, A.N., Shuvalova, L.D., Zubkova, O.A., Khomyakova, E.A. and Glushankova, L.N., 2021. STIM2 mediates excessive store-operated calcium entry in patient-specific iPSC-derived neurons modeling a juvenile form of huntington’s disease. Frontiers in Cell and Developmental Biology, 9, p.625231. doi: 10.3389/fcell.2021.625231.

36. Saitsu, H., Fukai, R., Ben-Zeev, B., Sakai, Y., Mimaki, M., Okamoto, N., Suzuki, Y., Monden, Y., Saito, H., Tziperman, B. and Torio, M., 2016. Phenotypic spectrum of GNAO1 variants: epileptic encephalopathy to involuntary movements with severe developmental delay. European Journal of Human Genetics, 24(1), pp.129–134. doi: 10.1038/ejhg.2015.92.

37. Danti, F.R., Galosi, S., Romani, M., Montomoli, M., Carss, K.J., Raymond, F.L., Parrini, E., Bianchini, C., McShane, T., Dale, R.C. and Mohammad, S.S., 2017. GNAO1 encephalopathy: broadening the phenotype and evaluating treatment and outcome. Neurology Genetics, 3(2). doi: 10.1212/NXG.0000000000000143.

38. Knight, K.M., Ghosh, S., Campbell, S.L., Lefevre, T.J., Olsen, R.H., Smrcka, A.V., Valentin, N.H., Yin, G., Vaidehi, N. and Dohlman, H.G., 2021. A universal allosteric mechanism for G protein activation. Molecular cell, 81(7), pp.1384–1396. doi: 10.1016/j.molcel.2021.02.002.

39. Novelli, M., Galosi, S., Zorzi, G., Martinelli, S., Capuano, A., Nardecchia, F., Granata, T., Pollini, L., Di Rocco, M., Marras, C.E. and Nardocci, N., 2023. GNAO1-related movement disorder: An update on phenomenology, clinical course, and response to treatments. Parkinsonism & Related Disorders, p.105405. doi: 10.1016/j.parkreldis.2023.105405.

40. Jeon, S.J., Kim, J.W., Kim, K.C., Han, S.M., Go, H.S., Seo, J.E., Choi, C.S., Ryu, J.H., Shin, C.Y. and Song, M.R., 2014. Translational regulation of NeuroD1 expression by FMRP: involvement in glutamatergic neuronal differentiation of cultured rat primary neural progenitor cells. Cellular and molecular neurobiology, 34, pp.297–305. doi: 10.1007/s10571-013-0014-9.

41. Callan, M.A., Cabernard, C., Heck, J., Luois, S., Doe, C.Q. and Zarnescu, D.C., 2010. Fragile X protein controls neural stem cell proliferation in the Drosophila brain. Human molecular genetics, 19(15), pp.3068–3079. doi: 10.1093/hmg/ddq213.

42. Magri, L., Cambiaghi, M., Cominelli, M., Alfaro-Cervello, C., Cursi, M., Pala, M., Bulfone, A., Garcìa-Verdugo, J.M., Leocani, L., Minicucci, F. and Poliani, P.L., 2011. Sustained activation of mTOR pathway in embryonic neural stem cells leads to development of tuberous sclerosis complex-associated lesions. Cell stem cell, 9(5), pp.447–462. DOI 10.1016/j.stem.2011.09.008

43. Valli, E., Trazzi, S., Fuchs, C., Erriquez, D., Bartesaghi, R., Perini, G. and Ciani, E., 2012. CDKL5, a novel MYCN-repressed gene, blocks cell cycle and promotes differentiation of neuronal cells. Biochimica et Biophysica Acta (BBA)-Gene Regulatory Mechanisms, 1819(11-12), pp.1173–1185 10.1016/j.bbagrm.2012.08.001

44. Wong, L.C., Singh, S., Wang, H.P., Hsu, C.J., Hu, S.C. and Lee, W.T., 2019. FOXG1-related syndrome: from clinical to molecular genetics and pathogenic mechanisms. International Journal of Molecular Sciences, 20(17), p.4176. doi: 10.3390/ijms20174176.

45. Darbandi, S.F., Schwartz, S.E.R., Qi, Q., Catta-Preta, R., Pai, E.L.L., Mandell, J.D., Everitt, A., Rubin, A., Krasnoff, R.A., Katzman, S. and Tastad, D., 2018. Neonatal Tbr1 dosage controls cortical layer 6 connectivity. Neuron, 100(4), pp.831–845. doi: 10.1016/j.neuron.2018.09.027

46. Crespo, I., Pignatelli, J., Kinare, V., Méndez-Gómez, H.R., Esgleas, M., Román, M.J., Canals, J.M., Tole, S. and Vicario, C., 2022. Tbr1 misexpression alters neuronal development in the cerebral cortex. Molecular Neurobiology, 59(9), pp.5750–5765. doi: 10.1007/s12035-022-02936-x.

47. Mendez-Gomez, H.R., Vergaño-Vera, E., Abad, J.L., Bulfone, A., Moratalla, R., de Pablo, F. and Vicario-Abejón, C., 2011. The T-box brain 1 (Tbr1) transcription factor inhibits astrocyte formation in the olfactory bulb and regulates neural stem cell fate. Molecular and Cellular Neuroscience, 46(1), pp.108–121. 10.1016/j.mcn.2010.08.011

48. Pacey, L.K., Guan, S., Tharmalingam, S., Thomsen, C. and Hampson, D.R., 2015. Persistent astrocyte activation in the fragile X mouse cerebellum. Brain and behavior, 5(10), p.e00400. doi: 10.1002/brb3.400.

49. Sunamura, N., Iwashita, S., Enomoto, K., Kadoshima, T. and Isono, F., 2018. Loss of the fragile X mental retardation protein causes aberrant differentiation in human neural progenitor cells. Scientific Reports, 8(1), p.11585. doi: 10.1038/s41598-018-30025-4.

50. Chatterjee, P., Pedrini, S., Stoops, E., Goozee, K., Villemagne, V.L., Asih, P.R., Verberk, I.M., Dave, P., Taddei, K., Sohrabi, H.R. and Zetterberg, H., 2021. Plasma glial fibrillary acidic protein is elevated in cognitively normal older adults at risk of Alzheimer’s disease. Translational psychiatry, 11(1), p.27. doi: 10.1038/s41398-020-01137-1.

51. Ganne, A., Balasubramaniam, M., Griffin, W.S.T., Shmookler Reis, R.J. and Ayyadevara, S., 2022. Glial fibrillary acidic protein: a biomarker and drug target for alzheimer’s disease. Pharmaceutics, 14(7), p.1354. doi: 10.3390/pharmaceutics14071354.

52. Sloan, S.A. and Barres, B.A., 2014. Mechanisms of astrocyte development and their contributions to neurodevelopmental disorders. Current opinion in neurobiology, 27, pp.75–81. 10.1016/j.conb.2014.03.005

53. Escartin, C., Galea, E., Lakatos, A., O’Callaghan, J.P., Petzold, G.C., Serrano-Pozo, A., Steinhäuser, C., Volterra, A., Carmignoto, G., Agarwal, A. and Allen, N.J., 2021. Reactive astrocyte nomenclature, definitions, and future directions. Nature neuroscience, 24(3), pp.312–325. doi: 10.1038/s41593-020-00783-4.

54. Verhoog, Q.P., Holtman, L., Aronica, E. and van Vliet, E.A., 2020. Astrocytes as guardians of neuronal excitability: mechanisms underlying epileptogenesis. Frontiers in Neurology, 11, p.591690. 10.3389/fneur.2020.591690

55. Steinberg, D.J., Repudi, S., Saleem, A., Kustanovich, I., Viukov, S., Abudiab, B., Banne, E., Mahajnah, M., Hanna, J.H., Stern, S. and Carlen, P.L., 2021. Modeling genetic epileptic encephalopathies using brain organoids. EMBO Molecular Medicine, 13(8), p.e13610. 10.15252/emmm.202013610

56. Thompson, J.A., Miralles, R.M., Wengert, E.R., Wagley, P.K., Yu, W., Wenker, I.C. and Patel, M.K., 2022. Astrocyte reactivity in a mouse model of SCN8A epileptic encephalopathy. Epilepsia Open, 7(2), pp.280–292. 10.1002/epi4.12564

57. Uhlmann, E.J., Wong, M., Baldwin, R.L., Bajenaru, M.L., Onda, H., Kwiatkowski, D.J., Yamada, K. and Gutmann, D.H., 2002. Astrocyte-specific TSC1 conditional knockout mice exhibit abnormal neuronal organization and seizures. Annals of neurology, 52(3), pp.285–296. 10.1002/ana.10283

58. Çarçak, N., Onat, F. and Sitnikova, E., 2023. Astrocytes as a target for therapeutic strategies in epilepsy: current insights. Frontiers in Molecular Neuroscience, 16.10.3389/fnmol.2023.1183775

59. Kim, Y.T., Hur, E.M., Snider, W.D. and Zhou, F.Q., 2011. Role of GSK3 signaling in neuronal morphogenesis. Frontiers in molecular neuroscience, 4, p.48. doi: 10.3389/fnmol.2011.00048.

60. González, M.I. and Brooks-Kayal, A., 2011. Altered GABAA receptor expression during epileptogenesis. Neuroscience letters, 497(3), pp.218–222. 10.1016/j.neulet.2011.02.052

61. Lunev, E.A., Shmidt, A.A., Vassilieva, S.G., Savchenko, I.M., Loginov, V.A., Marina, V.I., Egorova, T.V. and Bardina, M.V., 2022. Effective viral delivery of genetic constructs to neuronal culture for modeling and Gene therapy of Gnao1 encephalopathy. Molecular Biology, 56(4), pp.559–571. doi: 10.31857/S0026898422040061.

62. Shuvalova, L.D., Eremeev, A.V., Bogomazova, A.N., Novosadova, E.V., Zerkalenkova, E.A., Olshanskaya, Y.V., Fedotova, E.Y., Glagoleva, E.S., Illarioshkin, S.N., Lebedeva, O.S. and Lagarkova, M.A., 2020. Generation of induced pluripotent stem cell line RCPCMi004-A derived from patient with Parkinson’s disease with deletion of the exon 2 in PARK2 gene. Stem Cell Research, 44, p.101733. doi: 10.1016/j.scr.2020.101733.

63. Whye, D., Wood, D., Saber, W.A., Norabuena, E.M., Makhortova, N.R., Sahin, M. and Buttermore, E.D., 2023. A Robust Pipeline for the Multi-Stage Accelerated Differentiation of Functional 3D Cortical Organoids from Human Pluripotent Stem Cells. Current Protocols, 3(1), p.e641. doi: 10.1002/cpz1.641.

64. Langmead, B. and Salzberg, S.L., 2012. Fast gapped-read alignment with Bowtie 2. Nature methods, 9(4), pp.357–359. doi: 10.1038/nmeth.1923.

65. Dobin, A., Davis, C.A., Schlesinger, F., Drenkow, J., Zaleski, C., Jha, S., Batut, P., Chaisson, M. and Gingeras, T.R., 2013. STAR: ultrafast universal RNA-seq aligner. Bioinformatics, 29(1), pp.15–21. doi: 10.1093/bioinformatics/bts635.

66. Harrow, J., Frankish, A., Gonzalez, J.M., Tapanari, E., Diekhans, M., Kokocinski, F., Aken, B.L., Barrell, D., Zadissa, A., Searle, S. and Barnes, I., 2012. GENCODE: the reference human genome annotation for The ENCODE Project. Genome research, 22(9), pp.1760–1774. doi: 10.1101/gr.135350.111.

67. Love, M.I., Huber, W. and Anders, S., 2014. Moderated estimation of fold change and dispersion for RNA-seq data with DESeq2. Genome biology, 15(12), pp.1–21. doi: 10.1186/s13059-014-0550-8.

68. Zhu, A., Ibrahim, J.G. and Love, M.I., 2019. Heavy-tailed prior distributions for sequence count data: removing the noise and preserving large differences. Bioinformatics, 35(12), pp.2084–2092. doi: 10.1093/bioinformatics/bty895.

69. Conway, J.R., Lex, A. and Gehlenborg, N., 2017. UpSetR: an R package for the visualization of intersecting sets and their properties. Bioinformatics, 33(18), pp.2938–2940. doi: 10.1093/bioinformatics/btx364.

70. Gu, Z., Eils, R. and Schlesner, M., 2016. Complex heatmaps reveal patterns and correlations in multidimensional genomic data. Bioinformatics, 32(18), pp.2847–2849. doi: 10.1093/bioinformatics/btw313.

71. Liao, Y., Wang, J., Jaehnig, E.J., Shi, Z. and Zhang, B., 2019. WebGestalt 2019: gene set analysis toolkit with revamped UIs and APIs. Nucleic acids research, 47(W1), pp.W199–W205. doi: 10.1093/nar/gkz401.

