## Supplementary material for "Cortical neurons obtained from patient-derived iPSCs with GNAO1 p.G203R variant show altered differentiation and functional properties"

A

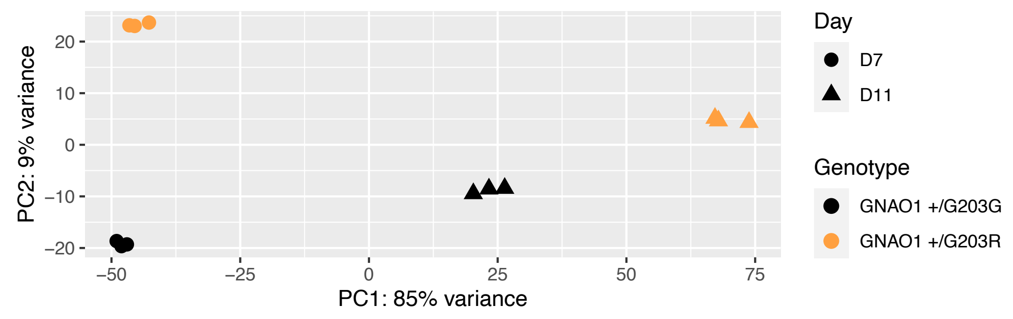


B

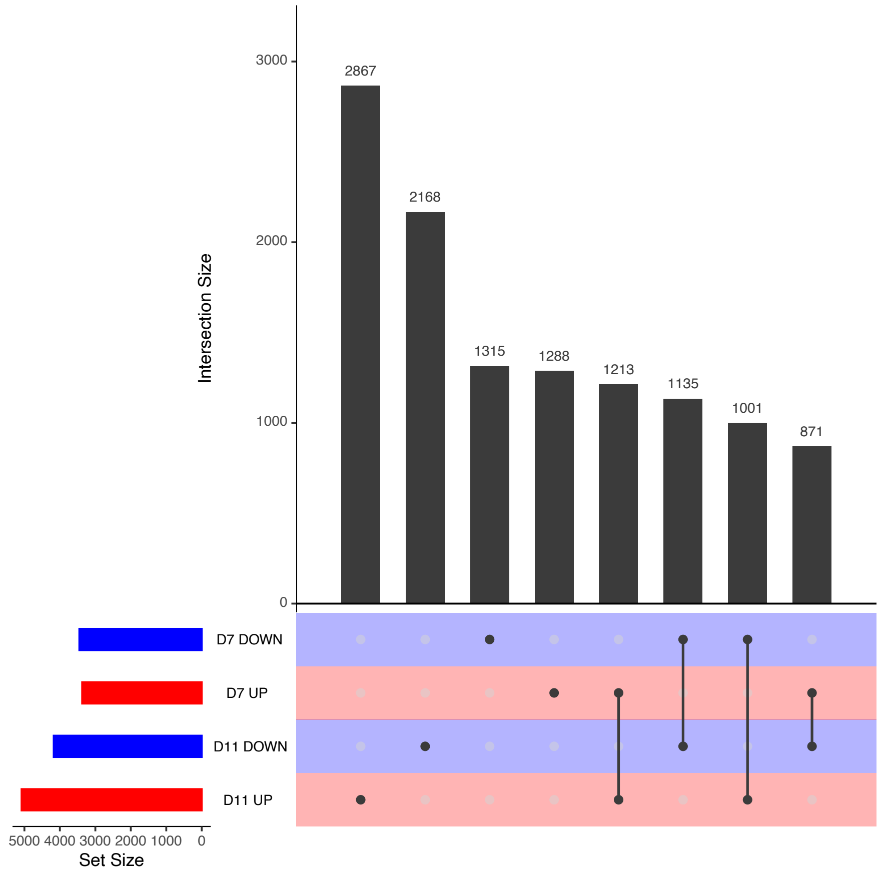


**Supplementary Figure S1. RNA-Seq analysis: PCA and UpSet plot**

**A.** PCA plot of RNA-Seq data from GNAO1$^{+/G203R}$ and GNAO1$^{+/G203G}$ NPCs at day 7 and 11. **B.** UpSet plot showing the overlap of genes that are differentially expressed between GNAO1$^{+/G203R}$ and GNAO1$^{+/G203G}$ NPCs (adjusted *p*-value < 0.01) at the two time points.

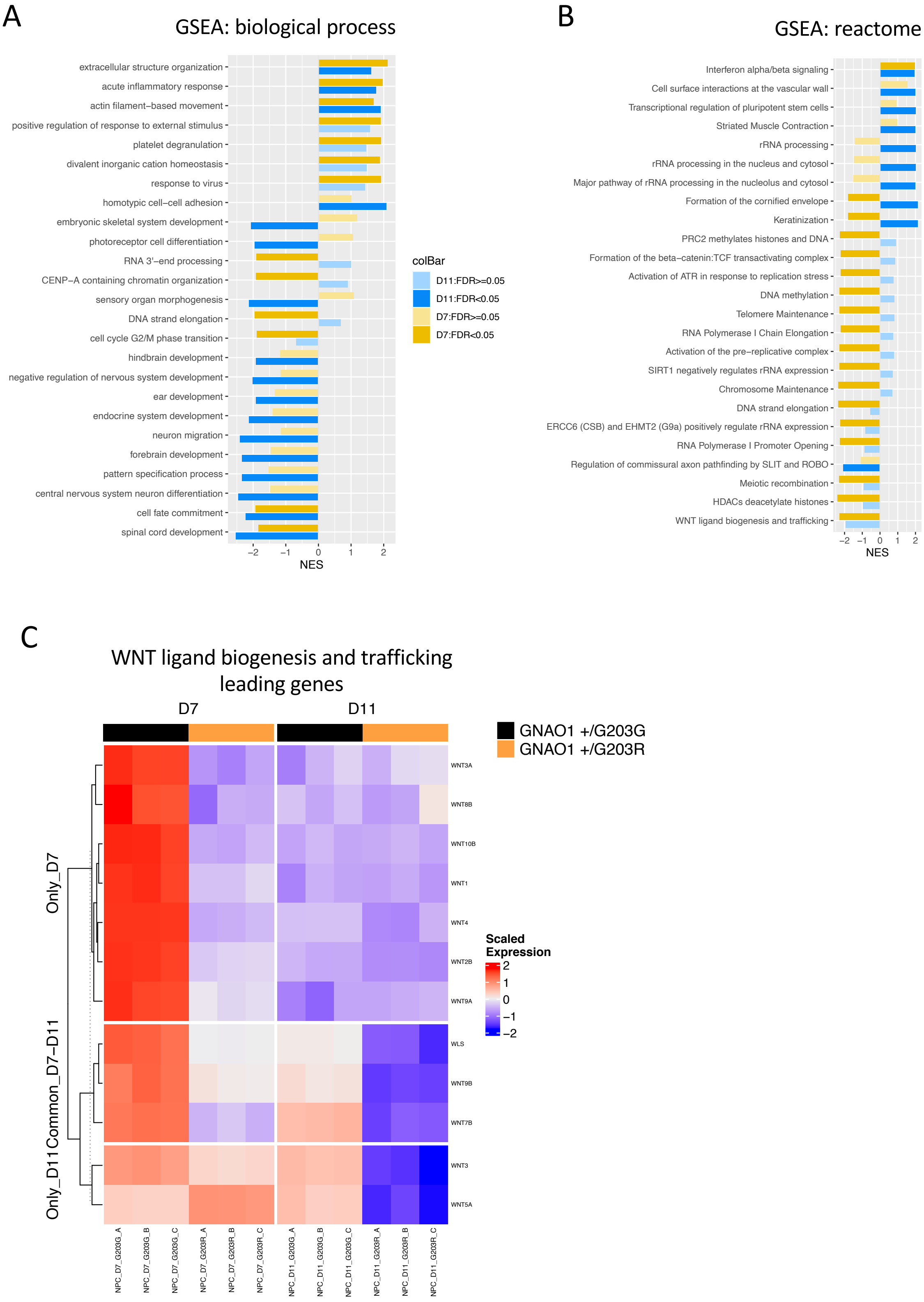
A
GSEA: biological process
extracellular structure organization
acute inflammatory response
actin filament−based movement
positive regulation of response to external stimulus
platelet degranulation
divalent inorganic cation homeostasis
response to virus
homotypic cell−cell adhesion
embryonic skeletal system development
photoreceptor cell differentiation
RNA 3'−end processing
CENP−A containing chromatin organization
sensory organ morphogenesis
DNA strand elongation
cell cycle G2/M phase transition
hindbrain development
negative regulation of nervous system development
ear development
endocrine system development
neuron migration
forebrain development
pattern specification process
central nervous system neuron differentiation
cell fate commitment
spinal cord development
NES
colBar
D11:FDR>=0.05
D11:FDR<0.05
D7:FDR>=0.05
D7:FDR<0.05
B
GSEA: reactome
Interferon alpha/beta signaling
Cell surface interactions at the vascular wall
Transcriptional regulation of pluripotent stem cells
Striated Muscle Contraction
rRNA processing
rRNA processing in the nucleus and cytosol
Major pathway of rRNA processing in the nucleolus and cytosol
Formation of the cornified envelope
Keratinization
PRC2 methylates histones and DNA
Formation of the beta−catenin:TCF transactivating complex
Activation of ATR in response to replication stress
DNA methylation
Telomere Maintenance
RNA Polymerase I Chain Elongation
Activation of the pre−replicative complex
SIRT1 negatively regulates rRNA expression
Chromosome Maintenance
DNA strand elongation
ERCC6 (CSB) and EHMT2 (G9a) positively regulate rRNA expression
RNA Polymerase I Promoter Opening
Regulation of commissural axon pathfinding by SLIT and ROBO
Meiotic recombination
HDACs deacetylate histones
WNT ligand biogenesis and trafficking
NES
C
WNT ligand biogenesis and trafficking leading genes
D7
D11
GNAO1 +/G203G
GNAO1 +/G203R
Only_D7
Common_D7−D11
Only_D11
WNT3A
WNT8B
WNT10B
WNT1
WNT4
WNT2B
WNT9A
WLS
WNT9B
WNT7B
WNT3
WNT5A
Scaled Expression
NPC_D7_G203G_A
NPC_D7_G203G_B
NPC_D7_G203G_C
NPC_D7_G203R_A
NPC_D7_G203R_B
NPC_D7_G203R_C
NPC_D11_G203G_A
NPC_D11_G203G_B
NPC_D11_G203G_C
NPC_D11_G203R_A
NPC_D11_G203R_B
NPC_D11_G203R_C

**Supplementary Figure S2. GSEA results**

**A-C.** Barplots showing the results of the GSEAs performed on the GNAO1$^{+/G203R}$ vs GNAO1$^{+/G203G}$ contrast at day 7 and day 11. Only categories with absolute Normalized Enrichment Score (NES) > 1.7 and FDR-adjusted p-value < 0.05 in at least one time point are reported. Functional categories are indicated above each panel. The Gene Ontology (GO) terms correspond to the non-redundant sets provided by WebGestalt. A positive or negative NES indicates that the gene set is enriched among the upregulated or the downregulated genes, respectively. **D.** Heatmap showing the expression in GNAO1$^{+/G203R}$ and GNAO1$^{+/G203G}$ NPCs at day 7 and 11 of the leading edge genes identified via GSEA for the "WNT ligand biogenesis and trafficking" Reactome pathway. Genes that were identified as leading edge ones only at day 7, only at day 11, or at both time points were split into three groups (Only_D7, Only D11, Common_D7-D11, respectively). The reported expression values correspond to row-scaled (Z-score), rlog-transformed count data.

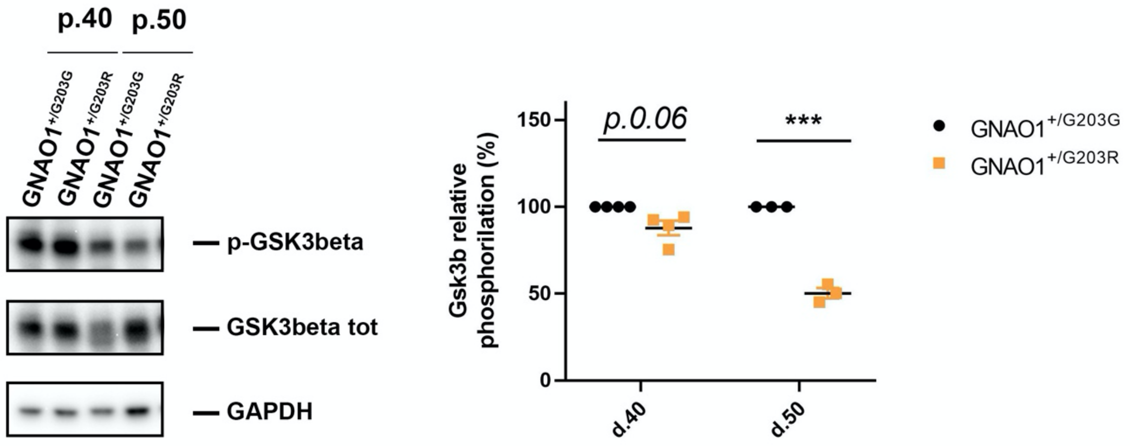


**Supplementary Figure S3. Phospho-GSK3β levels at days 40 and 50 of differentiation** Left panel, representative western blot analysis showing levels of phospho-Gsk3β, Gsk3β and GAPDH in differentiating GNAO1-G203R neurons and control isogenic cells (isogenic CTR), at day 40 and 50. Right panel, dot plot shows the quantification of Gsk3β phosphorylation levels in control and p.G203R neurons at day 40 and 50. Graphs indicate the % of phosphoproteins normalized to GAPDH, indicating the total amount of phosphorylated protein inside the cells. Average values and standard deviation are also shown (two tails paired Student's t test; ***$p<0.001$).

| Primer name | Primer sequence 5'-3' |
|---|---|
| ATP5O FW | ACTCGGGTTTGACCTACAGC |
| ATP5O RV | GGTACTGAAGCATCGCACCT |
| FOXG1 FW | AGGAGGGCGAGAAGAAGAAC |
| FOXG1 RV | TCACGAAGCACTTGTTGAGG |
| NESTIN FW | GCGTTGGAACAGAGGTTGGA |
| NESTIN RV | TGGGAGCAAAGATCCAAGAC |
| GFAP FW | GATCAACTCACCGCCAACAG |
| GFAP RV | ATAGGCAGCCAGGTTGTTCT |
| PAX6 FW | ATGTGTGAGTAAAATTCTGGGCA |
| PAX6 RV | GCTTACAACTTCTGGAGTCGCTA |
| TBR2 FW | CTTCTTCCCGGAGCCCTTTGTC |
| TBR2 RV | TTCGCTCTGTTGGGGTGAAAGG |
| TBR1 FW | CAACGGAGCCTACAACAGCCTC |
| TBR1 RV | TGGTAGAACGGAGCTCCTTGGT |
| OTX2 FW | CAAAGTGAGACCTGCCAAAAAGA |
| OTX2 RV | TGGACAAGGGATCTGACAGTG |

**Supplementary Table S2. Sequences of the primers used in this study**
